# A nanoparticle priming agent reduces cellular uptake of cell-free DNA and enhances the sensitivity of liquid biopsies

**DOI:** 10.1101/2023.01.13.524003

**Authors:** Carmen Martin-Alonso, Shervin Tabrizi, Kan Xiong, Timothy Blewett, Sahil Patel, Zhenyi An, Sainetra Sridhar, Ahmet Bekdemir, Douglas Shea, Ava P. Amini, Shih-Ting Wang, Jesse Kirkpatrick, Justin Rhoades, Todd R. Golub, J. Christopher Love, Viktor A. Adalsteinsson, Sangeeta N. Bhatia

**Affiliations:** Koch Institute for Integrative Cancer Research, Massachusetts Institute of Technology; Cambridge, MA, USA; Harvard-MIT Division of Health Sciences and Technology, Institute for Medical Engineering and Science, Massachusetts Institute of Technology; Cambridge, MA, USA; Broad Institute of MIT and Harvard; Cambridge, MA, USA; Department of Radiation Oncology, Massachusetts General Hospital; Boston, MA, USA; Harvard Medical School; Boston, MA, USA; Division of Pulmonary and Critical Care, Department of Medicine, Massachusetts General Hospital; Boston, MA, USA; Microsoft Research New England; Cambridge, MA, USA; Department of Chemical Engineering, Massachusetts Institute of Technology; Cambridge, MA, USA; Department of Electrical Engineering and Computer Science, Massachusetts Institute of Technology; Cambridge, MA, USA; Department of Medicine, Brigham and Women’s Hospital; Boston, MA, USA; Wyss Institute at Harvard University; Boston, USA; Howard Hughes Medical Institute; Cambridge, MA, USA

## Abstract

Liquid biopsies are enabling minimally invasive monitoring and molecular profiling of diseases across medicine, but their sensitivity remains limited by the scarcity of cell-free DNA (cfDNA) in blood. Here, we report an intravenous priming agent that is given prior to a blood draw to increase the abundance of cfDNA in circulation. Our priming agent consists of nanoparticles that act on the cells responsible for cfDNA clearance to slow down cfDNA uptake. In tumor-bearing mice, this agent increases the recovery of circulating tumor DNA (ctDNA) by up to 60-fold and improves the sensitivity of a ctDNA diagnostic assay from 0% to 75% at low tumor burden. We envision that this priming approach will significantly improve the performance of liquid biopsies across a wide range of clinical applications in oncology and beyond.

## Main

Liquid biopsy measurements, such as the analysis of cell-free DNA (cfDNA) in blood, enable non-invasive diagnosis, monitoring, and molecular profiling of human diseases (*1*). However, despite their rapidly growing use in prenatal testing (*2*), infectious diseases (*3*), oncology (*4*), and organ transplant monitoring (*5*) over the last two decades, the sensitivity of liquid biopsies continues to be inadequate for their broad incorporation into clinical practice. For instance, for oncology applications the detection of circulating tumor DNA (ctDNA) shed by cancer cells remains limited, especially in low disease burden settings (*6*). Namely, the sensitivity of liquid biopsy-based screening tests is low (~20% for Stage I cancer (*7*)); liquid biopsies can be inconclusive in up to 40% of patients with advanced cancer (*8*); and up to 75% of patients who test negative for minimal residual disease after surgery experience metastatic recurrence (*9*).

To improve the sensitivity of ctDNA liquid biopsies, most efforts thus far have focused *ex vivo* on sequencing (*10*) and analysis (*11*), such as tracking multiple somatic variants (*12–15*) and integrating orthogonal features like DNA methylation or fragmentation patterns (*16–18*). However, we posit that the greater barrier lies *in vivo* – in cfDNA biology itself. Most cfDNA is cleared from blood within minutes (*19*). This process leaves little to none of the low fraction of ctDNA (as low as 1-10 parts per million (*9*)) available for accurate detection from a blood draw, regardless of the performance of downstream detection assays. To sample more ctDNA, others have proposed drawing larger volumes of blood (*4*) or performing plasmapheresis (*20*), which involves extracorporeal processing of an individual’s full blood volume. However, drawing large volumes of blood may not always be possible or desirable, particularly in frail or ill patients, and plasmapheresis carries associated risks and requires expensive instrumentation. Alternatively, methods to sample more proximal to the tumor (*21*) or to increase the shedding of ctDNA have also been investigated (*22*, *23*). Yet, both approaches require *a priori* knowledge of the tumor site and more invasive procedures, making them suboptimal in the context of cancer screening or monitoring of presumed disease-free patients.

To overcome the scarcity of cfDNA in circulation and the limitations of previous approaches, here and in a companion paper (*ref #adf2332*) we describe the very first demonstrations of intravenous priming agents that are given 1-2 hours prior to a blood draw to transiently delay cfDNA clearance, and thus enable greater recovery of cfDNA. In this work, we report a nanoparticle priming agent that acts on the cells responsible for cfDNA clearance, those in the mononuclear-phagocyte system (MPS) of the liver and spleen (*24*), to slow down cfDNA uptake. Given that the majority of cfDNA circulates while bound to histone proteins as nanoparticulate mononucleosomes (~11nm in diameter) (*1*), we hypothesized that a competing nanoparticle that is efficiently uptaken by the MPS, such as a liposome, may attenuate the uptake of mononucleosomes and thus enhance the accumulation of cfDNA in blood (**Fig. 1**). Although the notion of saturating MPS uptake with a nanoparticle has been explored therapeutically to decrease the hepatic accumulation of nanomedicines (*25–29*), our work is, to the best of our knowledge, the first instance of employing this strategy to increase the abundance of an endogenous analyte for a diagnostic application. We demonstrate that a DSPE-based liposomal agent inhibits the uptake of mononucleosomes by macrophages *in vitro* and increases the recovery of endogenous cfDNA from a blood draw *in vivo*. We further assess the effects of our liposomal agent on ctDNA testing and observe improved detection of ctDNA molecules in two tumor models (up to 60-fold) and enhanced sensitivity of ctDNA testing for tumor detection (as significantly as from 0% to 75%). Taken together, these works introduce and demonstrate the potential utility of priming agents to improve the sensitivity of liquid biopsies with broad implications in oncology and beyond.

**Fig. 1.**
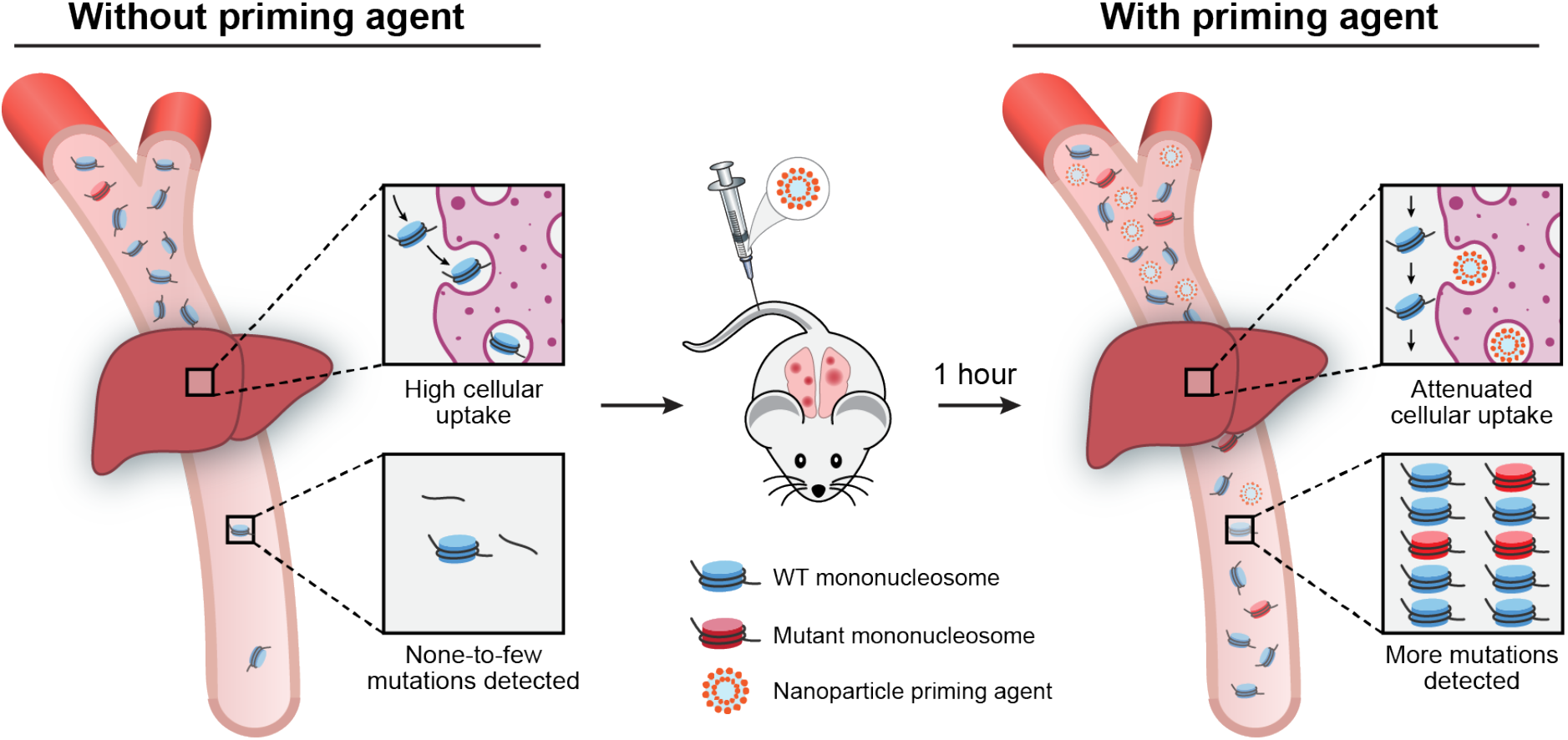
Schematic of the approach. In the absence of a priming agent, cfDNA is rapidly taken up by cells in the mononuclear-phagocyte system, yielding few molecules circulating for detection. Following intravenous administration of a nanoparticle priming agent, cellular uptake of nanoparticles attenuates uptake of cfDNA and results in more mutations being detected in blood.

To test our hypothesis that liposome treatment may inhibit macrophage uptake of cfDNA we first designed an *in vitro* 2D assay using the murine macrophage cell line J774A.1 (**Fig. 2A,B**). Following pre-treatment of J774A.1 cells with liposomes, we added Cy5-labeled mononucleosomes (**Fig. 2A, Fig. S1**) and quantified their uptake (**Fig. 2B**). To design a set of candidate priming agents, we prepared empty liposomes composed of cholesterol (50 mol%) and one of three different lipids, namely DSPE, DSPG, or DSPC (50 mol%), using the lipid film rehydration method. We selected these lipids as they are used in several FDA-approved liposome formulations (*30*) and were previously investigated in macrophage saturation studies (*26*, *27*, *29*). Given that the critical mass of the MPS resides in the liver, we designed the hydrodynamic diameter of the liposomes to be on par with the fenestrae of liver capillaries that measure 280nm in mice (*29*), such that nanoparticles would preferentially target liver-resident macrophages and minimize their interaction with hepatocytes. The three liposomal formulations exhibited comparable average diameters in the ~250nm range, but exhibited a range of surface charges as measured by dynamic light scattering (DLS; **Fig. 2C, Fig. S2**).

**Fig. 2.**
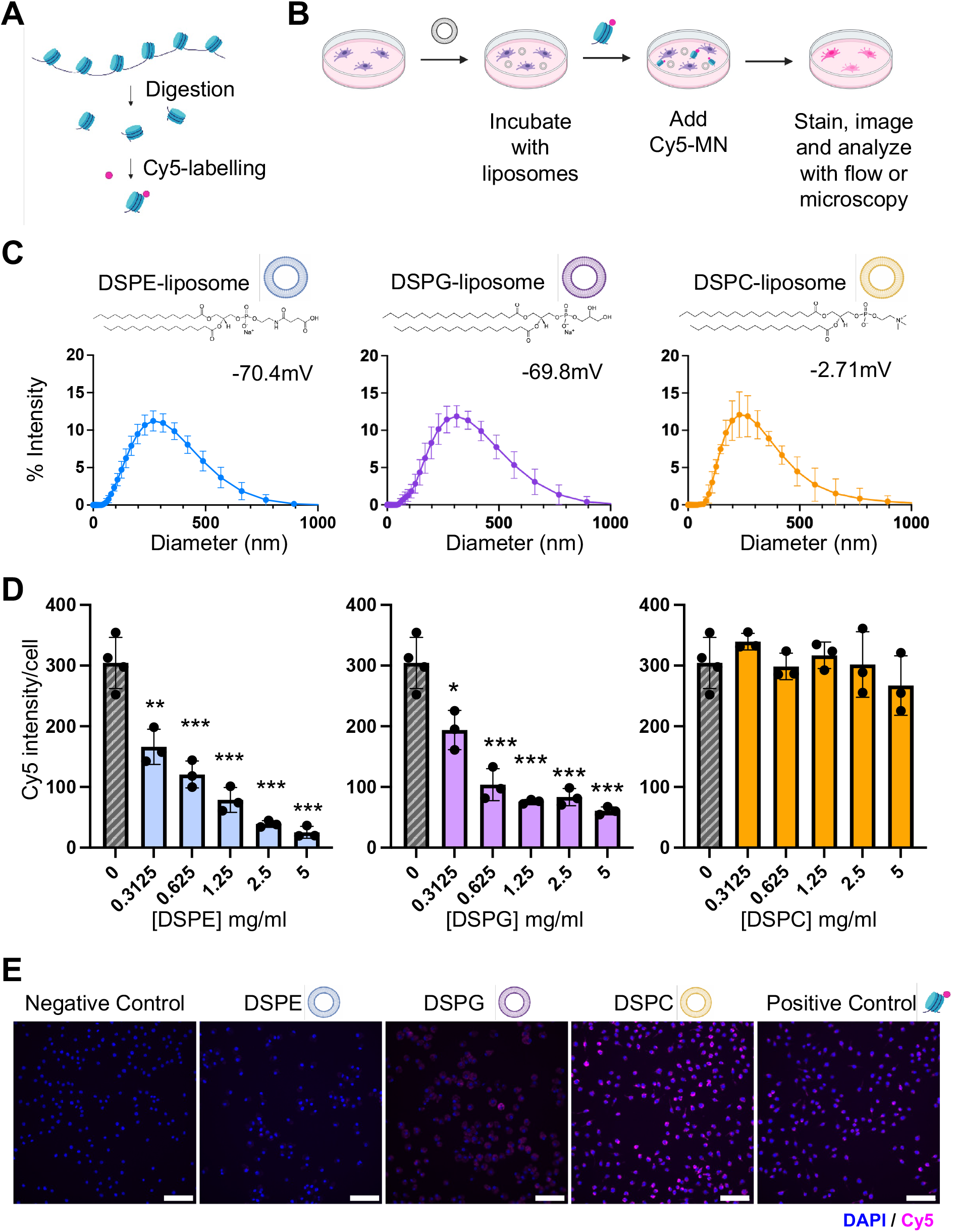
Select liposomal formulations inhibit the uptake of mononucleosomes by macrophages *in vitro*. Schematic of (**A**) preparation of Cy5-labelled mononucleosomes (Cy5-MN) from chromatin and (**B**) *in vitro* macrophage uptake inhibition assay. (**C**) Size distribution (average diameter ± s.d.: DSPE = 234nm ± 5.98 s.d., DSPG = 260nm ± 3.3 s.d. and DSPC = 227nm ± 3.3 s.d.) and zeta potential (DSPE = −70.4mV ± 1.32 s.d., DSPG = −69.8 mV± 1.0 s.d., and DSPC = −2.71mV ± 0.6 s.d.) of the three liposomal formulations, as measured via DLS. (**D**) Quantification of Cy5-MN uptake by J774A.1 cells after liposome pre-treatment for 4 hours as determined from epifluorescence images. Blue represents the DSPE-based, magenta the DSPG-based, and orange the DSPC-based liposome formulations (mean ± s.d., n = 3-4 wells per condition; * P < 0.05, ** P < 0.01, *** P < 0.001; oneway ANOVA). (**E**) Representative images of uptake of Cy5-MN from (D). From left to right, negative control (no liposomes or Cy5-MN), uptake following incubation with different liposomes at 5mg/ml, and positive control (no liposomes but Cy-MN), scale bars = 100μm.

The DSPE- and DSPG-based formulations significantly inhibited the uptake of mononucleosomes by macrophages in a dose-dependent manner (**Fig. 2D,E**; up to 12.0-fold and 5.0-fold for DSPE (P<0.0001) and DSPG (P<0.0001) liposomes respectively, relative to PBS control). However, the DSPC-based formulation had no effect on mononucleosome uptake even at the highest dose tested (Fig. 2d, 2e; *P* = 0.78). Given that the DSPE- and DSPG-based formulations are more negatively charged than the DSPC-based formulation, this trend is consistent with existing literature that has found that negatively charged particles display increased interactions with macrophages over those that are neutrally charged (*31*, *32*), possibly rendering them more potent agents for attenuating uptake by macrophages.

Using DSPE-liposomes, we confirmed independently that dose-dependent mononucleosome uptake inhibition was also observed in the macrophage cell line RAW264 (**Fig. S3A,B**) and that cell viability was not compromised with liposome treatment (**Fig. S3C,3D**). Additionally, we confirmed that liposomes did not impair phagocytosis in J774A.1 cells, suggesting that our priming agent would not affect phagocytic pathways that play an important role in defending the host from infection (*33*) (**Fig. S4**). Given the promising *in vitro* performance of the DSPE-based liposome, we selected this formulation for further *in vivo* characterization.

Having identified a nanoparticle formulation that inhibited mononucleosome uptake by macrophages *in vitro*, we next sought to determine whether liposome pre-treatment in healthy mice would inhibit the clearance of mononucleosomes *in vivo* and thus increase the accumulation of cfDNA in blood. To assess whether liposome administration affected cfDNA clearance, we pretreated healthy mice with liposomes and subsequently administered exogenous mononucleosomes carrying the Widom601 sequence (W601) (*34*) (**Fig. 3A**). Given that W601 does not align to the mouse genome, the abundance of mononucleosomes could be quantified longitudinally from plasma samples via qPCR. Consistent with decreased mononucleosome clearance upon liposome treatment, we observed that the percentage of W601 remaining in plasma 60 minutes after administration increased with an increasing dose of liposomes administered (**Fig. 3B**; median elevation between 100- and 1000-fold relative to PBS treatment at doses between 100 and 300mg/kg liposomes, P<0.05). These results were also in agreement with observed increases in the measured plasma half-life of mononucleosomes in mice pretreated with 100mg/kg and 300mg/kg liposomes, relative to mice treated with PBS (**Fig. 3C**).

**Fig. 3.**
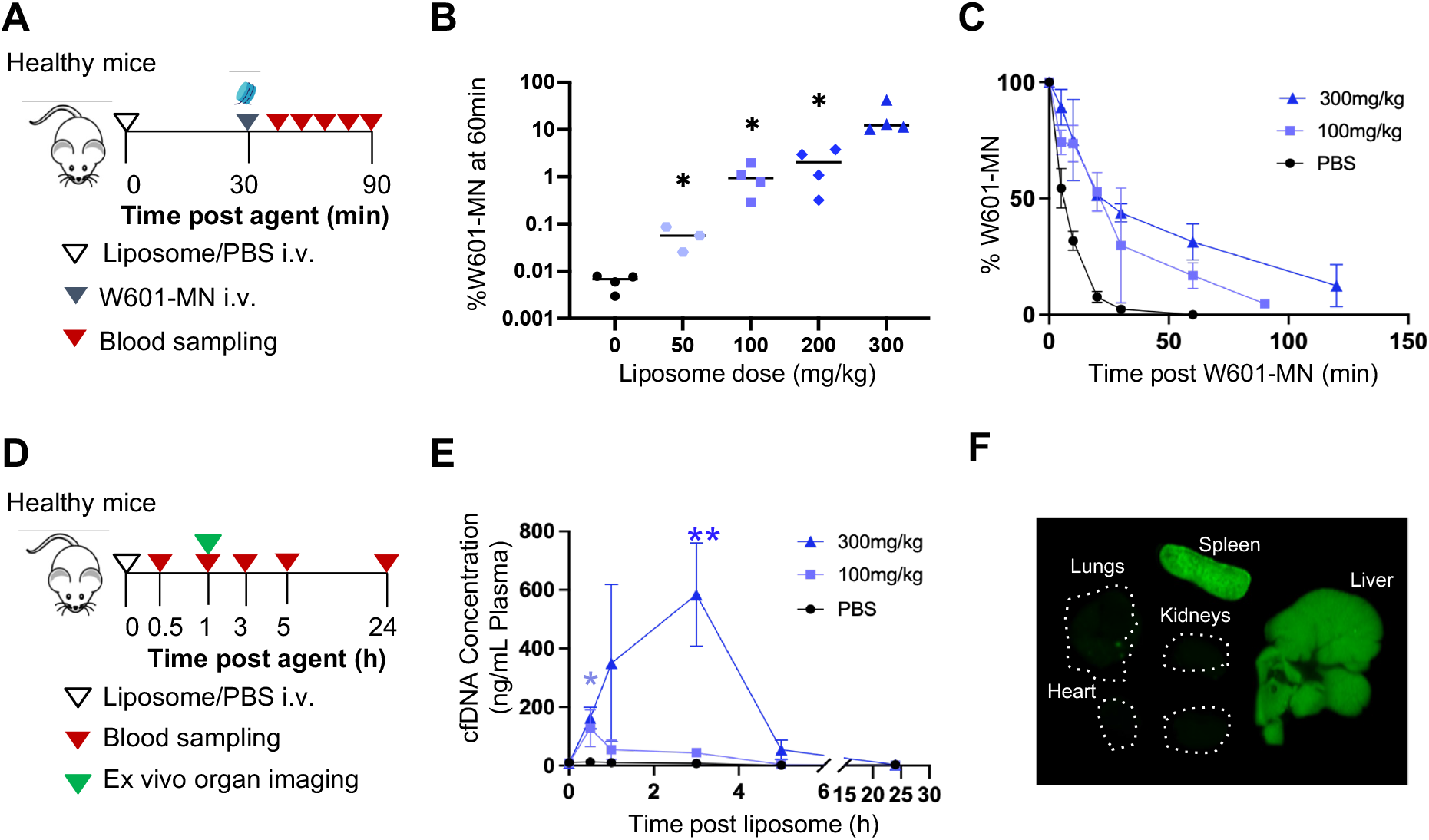
DSPE liposomes increase the recovery of cfDNA through decreased clearance in healthy mice. (**A**) Experimental timeline to determine the blood bioavailability and the half-life of W601-mononucleosomes (W601-MN) following liposome priming. Mice were treated with liposomes or PBS i.v. 30 min prior to i.v. administration of 1μg of W601-MN. Blood was sampled longitudinally, and plasma was analyzed to determine the percentage of W601-MN remaining in plasma from the time of injection. (**B**) Percentage of W601-MN remaining in plasma 60 min after administration of different liposome doses (median, *n* = 3-4 mice per group; * *P* < 0.05, two-tailed Mann-Whitney test). (**C**) Plasma clearance of W601-MN 30 min after the administration of PBS (t_1/2_ = 5.9 ± 0.7 min, 95% CI) or liposomes at 100 mg/kg (t_1/2_ = 19.6 ± 11.2 min) or 300 mg/kg (t_1/2_ = 24.9 ± 9.0 min) (mean ± s.d., *n* = 4 mice per group). (**D**) Experimental timeline to determine plasma cfDNA yields and liposome organ biodistribution. Mice were treated with either PBS, 100, or 300 mg/kg liposome, and plasma was collected longitudinally after injection to measure cfDNA yields. Additionally, 1 hour after Cy7-liposome administration, organs were harvested and imaged. (**E**) Plasma cfDNA yields following liposome administration, as measured via qPCR (mean ± s.d., n = 3 mice per group). Most significant elevations relative to PBS treatment with 100mg/kg treatment at 30 min (10.3-fold, \**P* = 0.034) and with 300mg/kg at 3 hours (78.0-fold, \*\**P* = 0.005); unpaired two-tailed t-test (n = 3 mice per group). (**F**) Organ biodistribution of Cy7-liposomes 1-hour after administration. Images from representative mouse shown (n = 4 mice). Dotted lines outline the perimeter of organs with low fluorescence intensity.

To determine whether decreased mononucleosome clearance would translate to higher endogenous plasma cfDNA levels, we administered liposomes or PBS control into healthy mice and measured cfDNA levels in blood collected longitudinally (**Fig. 3D**). Consistent with our hypothesis, the concentration of endogenous cfDNA in plasma spiked after liposome injection, with maximal recovery achieved 30 minutes after administration at the lower liposome dose tested (**Fig. 3E**; 10.3-fold increase over PBS, *P* = 0.034) or 3 hours after administration at the higher liposome dose (**Fig. 3E**; 78.0-fold increase over PBS, *P* = 0.005). Importantly, cfDNA levels returned to baseline within 5 hours or 24 hours of liposome treatment at the lower or higher dose, respectively (**Fig. 3E**), illustrating the transient effect of the nanoparticle agent on cfDNA recovery and supporting the potential use of such an agent for longitudinal monitoring applications.

We next set out to test whether MPS occupancy *in vivo* was driving the observed effect. To that end, we investigated how liver and spleen occupancy of liposomes related to the observed trends in cfDNA recovery over time by performing *in vivo* imaging of mice after the administration of Cy7-labeled liposomes (**Fig. S5**). We found that liposomes rapidly concentrated in target organs (**Fig. 3F, Fig. S5**), where they can interfere with cfDNA clearance. Together, these results suggest that *in vivo* the nanoparticle agent rapidly accumulates in MPS organs, decreases the plasma clearance of mononucleosomes, and increases the recovery of cfDNA from a blood draw.

Having demonstrated that liposome treatment increases the recovery of cfDNA in healthy mice, we next sought to determine whether our nanoparticle priming agent could improve the recovery of ctDNA. To this end, we turned to a flank tumor model of the colorectal cell line CT26, which was previously shown to shed DNA into circulation and thus deemed suitable for ctDNA testing (*35*). We inoculated CT26 tumors and treated mice with 700-1000mm^3^ burden with liposomes at different doses (ranging between 50mg/kg and 300mg/kg) or PBS control and collected blood prior to and 1 hour after treatment (**Fig. 4A**). We observed that cfDNA concentrations were significantly elevated in tumor-bearing mice 1 hour after liposome administration (*P* < 0.01) and further exhibited a dose-dependent increase relative to PBS-treated mice (**Fig. 4B**).

**Fig. 4.**
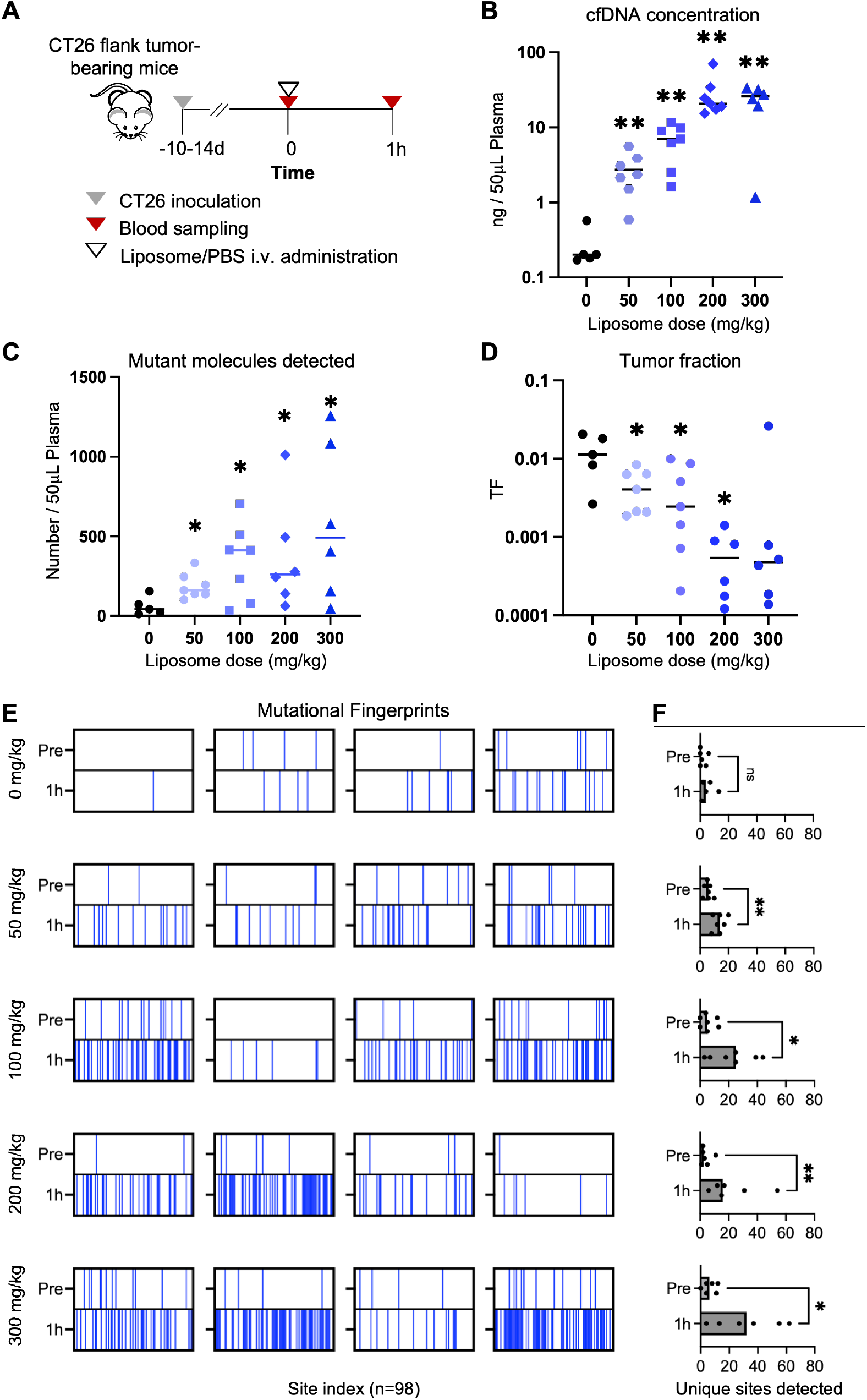
DSPE liposome priming increases the number of mutations detected in a flank tumor model. (**A**) Experimental timeline to detect mutations in the plasma of CT26 tumor-bearing mice using the nanoparticle priming agent. Blood was drawn prior to and 1 hour after i.v. administration of PBS or liposomes at a range of doses (0-300mg/kg). (**B**) cfDNA plasma concentration 1 hour after PBS or liposome treatment as quantified via qPCR (**C**) Number of mutant molecules (MT) per 50μl of plasma and (**D**) median tumor fractions detected 1 hour post treatment for each treatment group as quantified using the diagnostic ctDNA assay. Center line; median. (**E**) Mutational fingerprints showing individual sites detected for 4 representative mice per group. The top row represents mutations detected pre-treatment, and the bottom row represents mutations detected 1 hour posttreatment in a given plasma sample. Each vertical band corresponds to a different site in our 98-site mutational panel and is colored blue if detected at least once in the plasma sample. (**F**) Quantification of unique sites detected in (E). Bars; median. All panels refer to experiment with *n* = 5-7 mice per group. ns P > 0.05, * P < 0.05, ** P < 0.01, two-tailed Mann-Whitney test.

To enable tumor detection, we designed a ctDNA diagnostic assay tracking 98 mutations in the CT26 cell line (*36*) (**Fig. S6A**) based on an approach previously developed by our group (*12*). This assay utilizes duplex sequencing that requires a consensus from both strands of each DNA duplex to lower the sequencing error rate and can thus reliably detect single mutant DNA molecules and quantify tumor fractions as low as 1e-4 (**Fig. S6B**). Applying our ctDNA diagnostic assay to blood samples from the tumor-bearing mice (**Fig. S7 and Fig. S8, see Materials and Methods**), we observed that, as a result of increased cfDNA recovery, the recovery of mutant molecules was significantly enhanced following liposome treatment relative to PBS-treated mice (**Fig. 4C**; 3.8-fold, 9.7-fold, 6.1-fold, and 11.6-fold, *P* < 0.05 in mice treated with 50, 100, 200, and 300mg/kg liposomes, respectively). Moreover, the recovery of more mutant molecules after liposome treatment enabled the detection of additional unique mutational sites in our 98-probe panel relative to pre-treatment (**Fig. 4E,F, Fig. S9**; 2.3-fold, 5-fold, 6.4-fold, and 5.3-fold 1h after treatment with 50, 100, 200, and 300mg/kg liposomes, respectively). Importantly, the observed changes could not be explained by uneven tumor burden across groups, as we observed no statistically significant differences in pre-treatment plasma cfDNA concentration, mutant molecules, or tumor fractions between different treatment groups (**Fig. S10**).

Finally, while PBS treatment had a negligible impact on tumor fractions, liposome treatment significantly decreased tumor fractions, with tumor fractions trending towards lower values at higher doses (**Fig. 4D**). Given that white blood cells typically contribute the majority of cfDNA in circulation (*24*), we hypothesize that such dilutions in tumor fractions may be a result of increased cfDNA shedding by this cellular subset in response to high doses of liposomes, as has been previously reported (*37*). In addition to any safety concerns, dilutions in tumor fractions are also undesirable as they increase the cost of sequencing needed to come across any one particular mutant molecule.

The 100mg/kg liposome dose achieved comparable median fold-increase in mutant molecule recovery but resulted in less pronounced dilutions in tumor fractions relative to higher liposome doses. Additionally, at this dose we observed no significant changes in the absolute number of white blood cells 1 hour after administration relative to PBS treatment (**Fig. S11**, P > 0.05), and cfDNA yields returned to baseline shortly after administration (**Fig. 3F**). Given the performance and the favorable safety profile of the 100mg/kg liposome treatment, we selected this dose for assessing diagnostic ctDNA detection in tumor-bearing mice. Taken together, these results illustrate the utility of liposomes to both increase the likelihood to detect tumor-derived DNA molecules (**Fig. 4C**) as well as to improve the molecular profiling of tumors from cfDNA (**Fig. 4E,F**).

In humans, ctDNA levels typically correlate with tumor burden on imaging, making it easier to use ctDNA assays to detect later stage tumors (*38–40*). Despite the utility of the flank tumor model to identify an optimal liposome dose for ctDNA recovery enhancement, tumor fractions correlated poorly with tumor burden (**Fig. S12**), making it challenging to use this model to test whether liposomes can enable detection of smaller tumors. To overcome the limitation of this model, we tested the nanoparticle agent in a lung metastasis model that we found to have more physiologic shedding by virtue of its vascular seeding (**Fig. S12**).

In this model, the luciferized Luc-MC26 cell line was injected intravenously to establish lung metastases, and mice were treated once a week with 100mg/kg liposomes or PBS at different stages of tumor progression (**Fig. 5A**). A second ctDNA assay that tracked 1822-mutations in the Luc-MC26 cell-line was designed for ctDNA detection in this model (**Fig. S13**). We observed that treatment with the nanoparticle agent resulted in significantly increased plasma cfDNA concentration relative to PBS control (**Fig. 5B**; 7.1-fold, 14.0-fold, and 28.1-fold in weeks 1, 2, and 3, respectively) and significantly higher number of mutant molecules recovered relative to PBS control (**Fig. 5C**; 3.7-fold, 19.2-fold, and 59.5-fold at week 1, week 2, and week 3, respectively). As expected, mutant molecule recovery was enhanced across the entire range of tumor burdens analyzed (**Fig. 5D, Fig. S14**). Interestingly, we observed a stronger linear correlation between the number of mutant molecules detected and the tumor burden for liposome-treated mice (r = 0.820, *P* < 0.0001) than for PBS-treated mice (r = 0.128, *P* = 0.624) (**Fig. 5D**), which may be related to the stochasticity of mutant molecule detection in the PBS control group.

**Fig. 5.**
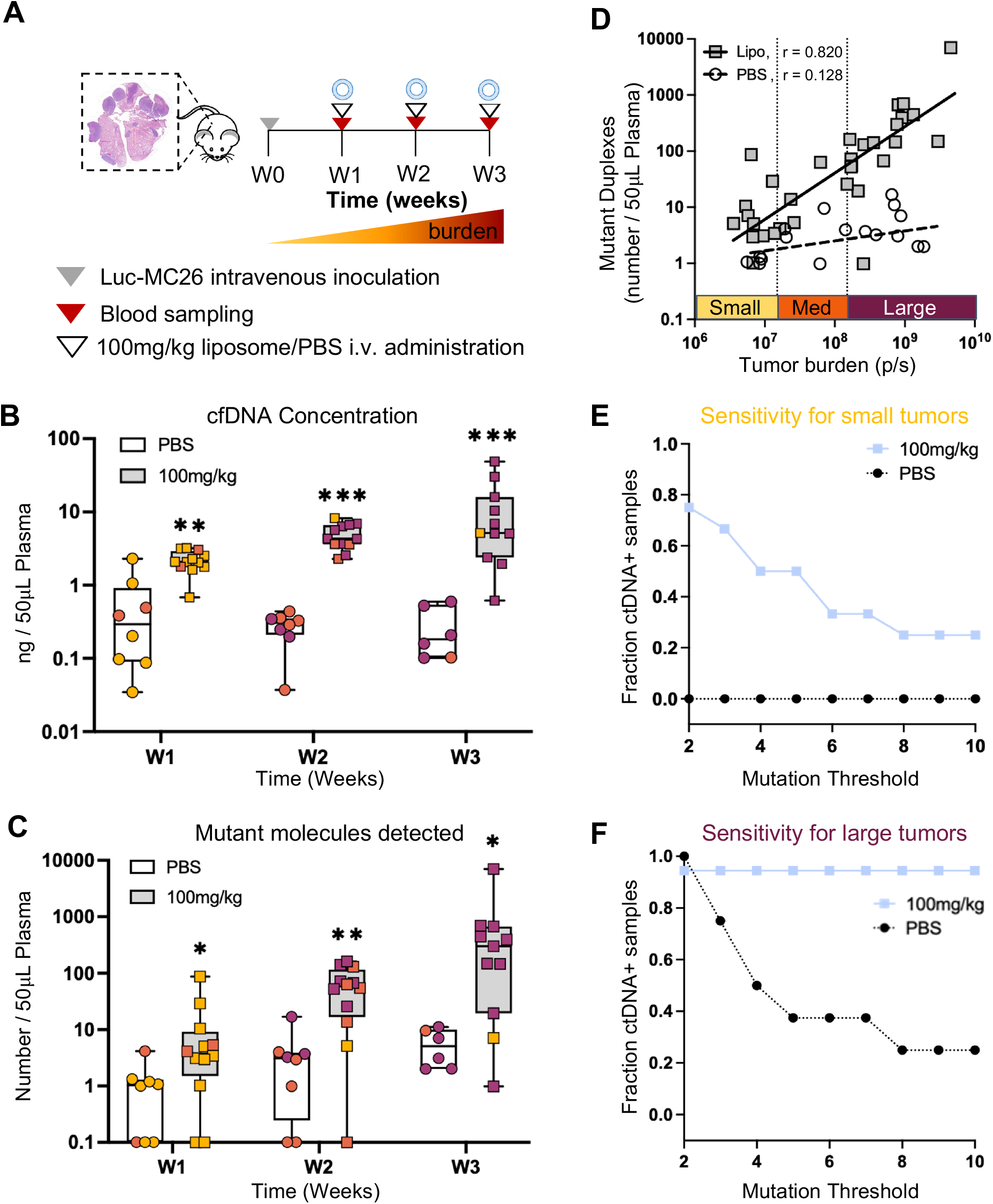
DSPE liposome priming increases the sensitivity of ctDNA testing in a CRC metastasis model and enables earlier detection. (**A**) Experimental timeline for the detection of mutations from the plasma of Luc-MC26 tumor-bearing mice. Blood was drawn prior to intravenous administration of PBS or liposomes (100mg/kg) and 1 hour after treatment 1 week, 2 weeks, and 3 weeks after tumor inoculation. (**B**) Plasma cfDNA yields and (**C**) number of mutant molecules detected in plasma 1 hour after PBS or liposome treatment at week 1,2, or 3. (Boxplot show median and interquartile range, *n* = 6-12 mice per group; * P < 0.05, ** P <0.01, *** P < 0.001, two-tailed Mann-Whitney test.). Yellow represents small tumors (burden < total flux 1.5e7 p/s), orange medium tumors (total flux 1.5e7 p/s < burden < total flux 1.5e8 p/s), and burgundy large tumors (burden > total flux 1.5e8 p/s). (**D**) Number of mutant molecules detected 1 hour after treatment vs. total tumor burden measured via *in vivo* bioluminescence. Pearson r for liposome-treated mice (r = 0.820, P < 0.0001) and for PBS-treated mice (r = 0.128, P = 0.624). (**E, F**) Sensitivity of ctDNA tests vs. mutation threshold for tumor detection for (E) low (burden < total flux 1.5e7 p/s) and (F) high (burden > total flux 1.5e8 p/s) tumor burden groups. Sensitivity was calculated as the fraction of samples for which the number of mutations detected in a blood sample after treatment was equal or exceeded a given mutation threshold.

To determine how the observed enhancement in mutant molecule recovery impacted the sensitivity of the ctDNA test, we classified each plasma sample as ctDNA positive if the number of mutations detected surpassed a given mutation threshold (between 2 and 10 mutations, from lower to higher stringency of the test). The nanoparticle agent improved the sensitivity of the ctDNA test, defined as the fraction of samples that were classified as ctDNA-positive, regardless of mutation threshold (**Fig. 5E,F**). The largest improvement in sensitivity was observed in the group with the lowest tumor burden (**Fig. 5D**). In particular, we observed that with a mutation threshold of 2, as has been previously applied for human samples (*12*), ctDNA was detected in 0% of low tumor burden mice injected with PBS compared to 75% of mice injected with liposomes (**Fig. 5E**). In mice with high tumor burden, the priming agent also proved useful by improving the robustness of the test and enabling application of a more stringent mutation threshold without compromising sensitivity (**Fig. 5F**).

Importantly, liposome treatment did not affect tumor progression (**Fig. S15**), and healthy mice on the same dosing schedule of liposomes at 100mg/kg or 300mg/kg showed no signs of toxicity, as determined by no significant changes in body weight over 28 days and no histological evidence of heart, lung, liver, spleen, or kidney toxicity 7 days after the last liposome injection **(Fig. S16**). Taken together, these results suggest that the nanoparticle agent can both increase the sensitivity and the reliability of ctDNA tests for tumor detection.

In conclusion, here we introduce the concept of a liquid biopsy priming agent that transiently increases the concentration of cfDNA in blood and enables sampling of more ctDNA for cancer detection. To achieve this, our priming agent attenuates the uptake capacity of cfDNA by the MPS system, increasing the amount of cfDNA recoverable from a blood draw. We first identified a DSPE-based liposomal agent that inhibits mononucleosome uptake *in vitro* and improves the recovery of cfDNA from blood *in vivo*. In two tumor models, we further demonstrated that this agent increases the recovery of ctDNA molecules by up to 60-fold, enhancing the sensitivity of a ctDNA diagnostic test at low tumor burden, the scenario most relevant for challenging early-stage or minimal-residual disease detection. Additionally, recovery of more of the tumor fingerprint following priming indicates that our agent may also yield a more robust test and improve other stages of cancer management like tumor genotyping for therapy selection or monitoring for drug resistant disease.

In this manner, our priming strategy could help overcome the fundamental limitation that most blood samples contain ultra-low levels of cfDNA (*41*, *42*), which today results in liquid biopsies with sensitivity that is too low to inform clinical decision making in many settings (*4*, *41*). Importantly, our ctDNA augmentation method should generalize well across all shedding tumors, as its mechanism of action is tumor-independent and does not require *in situ* exposure to external stimuli. Moreover, much like imaging contrast agents are routinely used for imaging scans, we believe that our nanoparticle agent will be safer and easier to incorporate into clinical practice relative to existing approaches to sample more cfDNA via collection of prohibitively large blood volumes or complex plasmapheresis procedures.

Although our work provides a proof of concept that attenuating cellular uptake of cfDNA can decrease its clearance from circulation, the dose of liposomes required to elicit such an effect and the subsequent dilution in tumor fraction observed must be addressed prior to clinical translation. Further optimization of agents for lower dosing as well as mechanistic studies into the dilution in tumor fraction will be necessary to meet these needs. To this end, we believe that the explosion of nanoparticle formulations being FDA-approved and entering clinical trials following the COVID-19 pandemic sets the stage for a more streamlined development of a nanoparticle priming agent, and will enhance formulation optimization, regulatory amenability, and physician adoption.

In addition, although the use of an injected priming agent undermines the minimally invasive nature of liquid biopsies, we believe that its use would be justifiable if it affords the requisite sensitivity to inform key clinical decisions, as is true for intravenous gadolinium and iodinated contrast agents. Alternatively, less-invasive routes of administration such as oral delivery could also be investigated. Despite the short-lived effect *in vivo*, the favorable safety profile in mice, and *in vitro* data suggesting that macrophages may retain their ability to clear infections, the safety of our liposomal agent must be assessed in larger mammals prior to clinical studies. Finally, the performance of our liposomal agent must be investigated across a larger range of cancer types to characterize individual-to-individual variability and to understand heterogeneities across cancer subtypes to support in-human clinical trials.

Our priming strategy should be of interest for oncology and beyond. Within oncology applications, this priming agent should also improve the performance of ctDNA analytical techniques beyond mutational profiling (*17*, *18*) and may even increase the recovery of cfDNA from other body fluids beyond plasma (*21*). Moreover, although we have limited our study to ctDNA testing in plasma for cancer detection, we believe that this work lays the groundwork for the potential development of priming agents for liquid biopsies at large. Given that our priming agent modulates cfDNA clearance, its use could be investigated beyond applications in oncology, such as for detection of microbial cfDNA during early infection (*3*) or for low abundance cfDNA biomarkers in other disease areas (*43–45*). Additionally, we believe that the concept of a priming agent that perturbs endogenous biomarker clearance *in vivo* marks a paradigm shift in how we think about the limit of detection of molecular diagnostics. The fact that our nanoparticle agent modulates *in vivo* cfDNA clearance to improve diagnostic performance opens the door to its use for other circulating biomarkers that are also cleared by the liver. In parallel, its distinct physicochemical characteristics to cfDNA particles could inspire the engineering of completely orthogonal priming agents for additional endogenous analytes.

In summary, here we present a first-in-class liquid biopsy priming agent capable of improving the sensitivity and the robustness of ctDNA testing in tumor-bearing mice by modulating cellular cfDNA clearance. Just as intravenous contrast agents have had profound impact on clinical imaging, we envision that molecular priming agents have the potential to expand the utility of liquid biopsies across all stages of cancer management and for applications beyond oncology.

## Materials and Methods

### Liposome synthesis

Liposomes were prepared using the lipid film re-hydration method with slight modifications from the protocol described by Saunders et al.(26) Briefly, ovine cholesterol (50 mol %, cat: 700000P, Avanti Polar Lipids) was solubilized in chloroform and added to 1,2-dipalmitoyl-sn-glycero-3-phosphoethanolamine-N-(succinyl) (sodium salt) (DSPE) (50 mol %, cat: 870225P, Avanti Polar Lipids), or 1,2-distearoyl-sn-glycero-3-phospho-(1’-rac-glycerol) (sodium salt) (DSPG) (50 mol %, cat: 840465P, Avanti Polar Lipids) or 1,2-distearoyl-sn-glycero-3-phosphocholine (DSPC) (50 mol %, cat: 850365P, Avanti Polar Lipids) with 1:1 (v/v) methanol. The solution was evaporated under nitrogen flow to form a thin dry film and vacuumed overnight to remove any traces of organic solvent. The lipid film was hydrated at 60oC with sterile DPBS to a total lipid concentration of 50mg/ml. Finally, extrusion was performed at 60oC with 1um (cat: WHA110410, MilliporeSigma) and 0.4um polycarbonate membranes (cat: WHA10417101, MilliporeSigma), 21 and 20 times respectively, using the 1000 ul Mini-Extruder from Avanti Polar Lipids (cat: 610023). For the fluorescent liposome used for biodistribution studies, 0.2 mol % of DSPE was replaced for Cy7-DSPE (cat: 810347C, Avanti Polar Lipids) prior to solubilization with organic solvents.

### Liposome characterization

The liposomes were characterized using a Zetasizer NanoZS (Malvern Instruments). To measure the hydrodynamic diameter and polydispersity index (PdI) of the liposomes, samples were diluted 1:100 in PBS and analyzed. To measure the zeta potential of the liposomes, 10ul of sample was added to 40ul of PBS and 850ul of deionized water. The morphology of liposomes was confirmed by cryogenic TEM (cryo-TEM) imaging. Prior to sample preparation, lacey copper grids coated with a continuous carbon film (LC200-Cu, Electron Microscopy Sciences) were plasma treated using a Denton Sputter Coater at 4 mA for 5 secs. The cryo-TEM samples were prepared on a Gatan Cryoplunge III at room temperature and 100% humidity. Liposome solution (3 μL, 5 mg/mL in PBS) was dropped on the plasma-treated grids and the excess of samples was removed by blotting for 4 sec before plunge freezing. The frozen grid was mounted on a Gatan 626 single-tilt cryo-transfer holder and imaged using a JEOL 2100F at 200 kV. All images were taken at 30 kx or 40 kx using the minimum dose exposure system and recorded on a Gatan UltraScan CCD camera.

### Mononucleosome extraction

To prepare mono-nucleosomes for labeling, chromatin was extracted from CT26 cells using the Nucleosome Preparation Kit (cat: 53504, Active Motif). The enzymatic digestion time for this cell line was optimized to 30min and the mononucleosome preparation protocol was followed as per manufacturer’s recommendations. To verify that chromatin had been successfully digested to mononucleosomes, the solution was subjected to DNA cleanup, and digestion efficiency was assessed via electrophoresis through a 1.5% agarose gel.

### Mononucleosome labeling

Aliquots of 10ug mononucleosomes were first washed and buffer-exchanged into PBS to remove any impurities from the preparation reagents. Four washes were performed using 30kDa Amicon filters (cat: UFC503024, EMD Millipore) by centrifugation at 12,000rpm, 10min, 4oC. After concentrating the volume of the last wash to 40ul, the protein yield was calculated using a commercial HeLa mononucleosome standard by measuring absorbance at 230nm using a Nanodrop 8000 Spectrophotometer (cat: ND-8000-GL, ThermoFisher). To label mononucleosomes, sulfonated-Cy5 (cat: 13320, Lumiprobe) was added at a 25 dye: 1 protein molar ratio, and the reaction incubated at 4oC in an Eppendorf Thermomixer C Model 5382 (Eppendorf) at 550rpm overnight. Excess dye was removed using Micro Bio-Spin^®^ Columns with Bio-Gel^®^ P-6 (cat:7326221, BioRad) by centrifugation at 1000g, 2min at room temperature. Labeling efficiency was measured by measuring Cy5 intensity at 650/680nm against a Cy5 standard using an Infinite F200 Pro reader (Tecan) fluorometer and protein yield by measuring absorbance at 230nm using a Nanodrop 8000 Spectrophotometer.

### Mononucleosome characterization

Mononucleosome stability post-labeling was confirmed by running a 4-12% Tris Glycine Novex gel with unlabeled or Cy5-labeled mononucleosomes (native running buffer, 4C, 150V, 90min). Colocalization of a Cy5-positive band with a protein band after Coomassie staining corresponding to the molecular weight of a mononucleosome was confirmed. Additionally, the morphology of mononucleosomes prepared from the chromatin of CT26 cells was compared to the morphology of mononucleosomes purchased commercially and used for in vivo pharmacokinetic studies (cat: 81070, ActiveMotif) using negative-stained TEM imaging. For sample preparation, first, ultrathin carbon film Au grids (CF300-Cu-UL, Electron Microscopy Sciences) were plasma treated using a Denton Sputter Coater at 5 mA for 8 secs. Next, 5 μL of NCP solution (17 ng/μL in PBS) was incubated on the plasma-treated grids for 1.5 mins and the excess of liquid was removed with a filter paper (pore size 110 nm, Whatman). The grid was washed twice by dropping 5 μL of deionized water on the grids and quickly removed using a filter paper. Negative staining was performed by dropping 5 μL of 2% uranyl acetate solution on the grid for 18–20 secs and the excess of liquid was removed by a filter paper. Imaging was performed on a JEOL 2100F TEM at 200 kV. All images were recorded on a Gatan UltraScan charge-coupled device (CCD) camera.

### Cell culture

For tumor inoculations, mouse cell lines CT26 (cat: CRL-2638, ATCC) and Luc-MC26 (carrying firefly luciferase, from the Kenneth K. Tanabe Laboratory, Massachusetts General Hospital) were cultured in RPMI-1640 (cat: R8758, Sigma) and ATCC-formulated Dulbecco’s Modified Eagle’s Medium DMEM (cat:10-013-CM, Corning), respectively. For in vitro mononucleosome uptake experiments, mouse macrophage cell lines RAW264 (TIB-71, ATCC) and J774A.1 (TIB-67, ATCC) were cultured in DMEM. Both media were supplemented with 10% fetal bovine serum (cat:100-500, GemCell) and 1% penicillin/streptomycin (cat: 30-002-C1, Corning), and cells were cultured in a humidified atmosphere of 95% air and 5% CO2 at 37oC. The CT26 and Luc-MC26 cell lines were declared pathogen free after being subjected to murine pathogen testing by the Diagnostic Laboratory of the Division of Comparative Medicine (MIT).

### In vitro macrophage mononucleosome uptake inhibition assay with liposomes

J774A.1 and RAW264 cells were plated at a density of 30,000 and 45,000 cells/chamber, respectively, in 8-well chamber slides (cat.80806, Ibidi). Following overnight acclimatization, cells were incubated with 300ul of liposomes (DSPE, DSPG, or DSPC) diluted in DMEM at the desired concentrations (0.1-5mg/ml) for 4h at 37oC. Next, 30ul of mononucleosomes were spiked into each well to achieve a final mononnucleosome concentration of 10nM and further incubated for 2h at 37oC. Cells incubated with DMEM followed by mononnucleosome addition were used as a positive control for uptake, and cells incubated only with DMEM were used as a negative control. At the end of the incubation, cells were washed once with DMEM, stained with Hoechst 33342 (cat: H3570, ThermoFisher) 1:2000 in DMEM for 10min, and further washed (twice with DMEM and once with PBS) to remove any extracellular mononnucleosomes. Subsequently, cells were fixed with 4% PFA for 20 min and washed with PBS prior to imaging on an Eclipse Ti microscope (Nikon).

To quantify cellular uptake, 4 fields of view per well were obtained at 10X mag and mean Cy5 fluorescence intensity per cell were quantified using custom scripts in QuPath (*46*). Results are displayed after background subtraction using the mean Cy5 fluorescence intensity per cell from the negative control. In parallel, quantification was also performed using flow cytometry by pooling cells from 2 wells per condition. To harvest cells for flow cytometry, cells were trypsinized with 50ul of trypsin/well (5min, 37oC) and quenched with 100ul of DMEM. Cells were scraped off the well and resuspended in 1ml of PBS. Cells were pelleted (350g, 5min, 25oC) and fixed with 100ul of 4%PFA (20min, 25oC). Cells were then washed with 1ml of FACS buffer (2% BSA in PBS) and resuspended in 400ul of FACS buffer prior to analysis in a LSRII-HTS flow cytometer (BD Biosciences). Excitation was performed with the 640nm laser and emission measured with the APC-A filter. Gating was performed to include only vertical and horizontal singlets. After gating, the percent positive cells was calculated as the fraction of cells above the APC value that marked the lower limit of the distribution for the positive control (treated with NCP only).

### In vitro cell viability and liposome uptake studies in macrophages

J774A.1 cells were seeded at 10,000 cells/well and RAW264 cells at 12,000 cells/well into two 96-well plates with transparent bottom. To measure liposome uptake, 24h after seeding, cells were incubated with Cy7-DSPE liposomes in DMEM at concentrations of 0-5mg/ml for 4 h and washed three times with PBS. Liposome uptake was then measured using an Infinite F200 Pro reader (Tecan) at 750/785nm. To measure cell viability, 24h after seeding, cells were incubated with DSPE liposomes at concentrations of 0-5mg/ml in DMEM containing the viability dye from the RealTime-Glo MT Cell Viability Assay (cat: G9712, Promega). After incubation with liposomes for 4h, endpoint cytotoxicity analysis was performed by measuring luminescence using an Infinite F200 Pro reader (Tecan) fluorometer. Percent viability was calculated relative to untreated cells in DMEM.

### In vitro E. coli uptake study in macrophages with liposomes

J774A.1 cells were seeded at 10,000 cells/well into 96-well plates (Greiner #655892). After 48 hours, cells were incubated with DSPE liposomes (50mg/ml) at 1:10, 1:20, and 1:50 dilution conditions for 1 hour in a cell culture incubator at 37oC with 5% CO2. Alexa 488 conjugated Escherichia coli (K-12 strain) BioParticles (cat: E13231, ThermoFisher) was added at 10ug/well. After 1h incubation at 37oC with 5% CO2, cells were fixed with 4% paraformaldehyde at room temperature for 20min and treated with 1x cell permeabilization buffer (cat: 00-8333-56, Invitrogen) for 10min. Cells were stained with Hoechst 33342 (1:5000, cat: H3570, ThermoFisher) and Phalloidin-Alexa 594 (1:2000, cat: 20553, Cayman) for 30min at room temperature. Cells were washed 5 times with 250ul PBS per well and hydrated with 50ul PBS per well. Images were taken using the Opera Phenix Imaging System (Perkin Elmer), and uptake of the E.coli bioparticles per cell was quantified automatically using the Harmony PhenoLOGIC software (Version 9.0, Perkin Elmer).

### Plasma collection

Blood was collected under anesthesia retro-orbitally for longitudinal studies and via terminal bleed at end point. For retro-orbital blood collections, 70ul of blood was collected from alternating eyes with non-heparinized hematocrit capillary tubes (cat: 22-362566, Fisher) and resuspended 1:1 (v/v) in 10mM EDTA in PBS (cat: 15575020, ThermoFisher). For terminal bleeds, mice were exsanguinated via cardiac puncture under anesthesia. Blood volume was measured, and 10mM EDTA in PBS was added 1:1 (v/v). All blood samples were kept on ice, and plasma was separated by centrifuging blood samples at 8,000 x g for 5 min at 4 °C within 2 hours of blood collection. Plasma was then stored at −80 °C until further use.

### Animal models

All animal studies were approved by the Massachusetts Institute of Technology Committee on Animal Care (MIT Protocol 0420-023-23). Animals were maintained in the Koch Institute animal facility with a 12h-light/12h-dark cycle at 18-23 °C and 50% humidity. All animals received humane care, and all experiments were conducted in compliance with institutional and national guidelines and supervised by staff from the Division of Comparative Medicine of the Massachusetts Institute of Technology. Female BALB/c mice (6-10 weeks, Taconic Biosciences) were used for NCP pharmacokinetic studies, liposome biodistribution studies, and assessment of the effects of liposomes on cfDNA yield over time. To generate the CT26 flank tumor model, female BALB/c mice (4-6 weeks, Taconic Biosciences) were injected subcutaneously with 2×106 CT26 cells resuspended in Opti-Mem (cat: 11058021, ThermoFisher) into bilateral rear flanks. Tumors were measured every other day for 2 weeks, and tumor volumes were calculated by the modified ellipsoidal formula: V = 0.5 * (l * w2), where l and w are the tumor length and width, respectively. To generate the lung metastasis model, 1×106 Luc-MC26 cells in 100ul DPBS were injected intravenously (i.v.) into female BALB/c mice (4-6 weeks, Taconic Biosciences). Tumor growth was monitored by luminescence using the In Vivo Imaging System (IVIS, PerkinElmer) on days 6, 13, and 20 after tumor inoculation.

### In vivo mononucleosome pharmacokinetic study

DSPE-liposomes or sterile DPBS were administered i.v. into awake mice (50-300mg/kg, 200ul). 30 min after liposome injection, 1ug recombinant mono-nucleosomes carrying the Widom601 (W601) sequence (cat: 81070, ActiveMotif) suspended in 10 uL DPBS were injected i.v. into anesthetized mice. In the study evaluating the % exoNCP remaining at 60min (n=4 per group), 35ul of blood was drawn retro-orbitally 1min and 60min after mononucleosome injection. % W601 remaining was calculated as the % W601 remaining at 60min relative to 1min, as quantified using Taqman qPCR (forward primer: 5’CGCTCAATTGGTCGTAGACA, reverse primer: 5’TATCTGACACGTGCCTGGAG and Taqman probe: /56-FAM/TC TAG CAC C/ZEN/G CTT AAA CGC ACG TA/3IABkFQ/). To obtain estimates of the mononucleosome half-life following liposome or PBS treatment (n=4 per group), 35ul of blood were collected from alternating eyes at 1, 5, 10, 20, 30, 60, 90, and 120 min and % W601 remaining was calculated relative to the W601 amount detected at 1min for samples over time. Half-life estimates were calculated by fitting an exponential decay equation model.

### Liposome biodistribution study

100mg/kg Cy7-DSPE-liposomes (200ul in sterile DPBS) were administered i.v. into awake mice. For ex vivo organ imaging, 1h after liposome administration mice were euthanized and liver, spleen, lungs, kidneys, and heart were harvested (n = 4 mice per group). A PBS-treated mouse was used as a negative control to measure organ autofluorescence. Organ fluorescence was measured using the 800nm filter of an Odyssey CLx instrument (Li-Cor). Biodistribution was quantified as the % of total fluorescence across all organs for each mouse. For in vivo liposome biodistribution studies (n = 1 mouse per group), 50-300mg/kg Cy7-liposomes (200ul in sterile DPBS) were administered i.v. in awake mice. Accumulation of liposomes in the liver and spleen was measured using the In Vivo Imaging System (IVIS, PerkinElmer) by defining regions of interest covering the upper abdomen of mice.

### Plasma cfDNA concentration measurements following liposome administration

100mg/kg or 300mg/kg DSPE-liposomes (200ul in sterile DPBS) or DPBS were administered i.v. in awake mice (n = 3 mice per group). 1min, 30min, 1h, 3h, 5h, and 24h after liposome administration, 70ul of blood was collected retro-orbitally. Only 2 blood samples were collected from each mouse, to prevent repeat sampling from the same capillary bed. Plasma cfDNA concentration was quantified as described in “cfDNA extraction and quantification”.

### Liposome dose-titration for tumor detection in the CT26 model

Mice bearing bilateral CT26-flank tumors were randomized into different treatment groups (300mg/kg, 200mg/kg, 100mg/kg, 50mg/kg DSPE-liposomes or PBS, n = 5-7 mice per group) at end point such that the tumor burden of the group ranged between 700-1000 mm3. As an internal control, 70ul blood was sampled retro-orbitally from each mouse prior to treatment. Subsequently, 50-300mg/kg DSPE-liposomes (in 200ul sterile DPBS) or sterile DPBS were administered i.v. into awake mice. 1h after treatment, 70ul of blood was collected retro-orbitally from the contralateral eye, and a terminal bleed was then performed, and blood collected for analysis. cfDNA concentration measurement and ctDNA detection was performed on all samples as described below.

### Sensitivity test of liposomal agent in the Luc-MC26 lung metastasis model

Six days after tumor inoculation, mice bearing Luc-MC26 metastatic tumors were randomized into different treatment groups (100mg/kg DSPE-liposomes (n = 12 mice) or PBS (n = 8 mice)) such that total burden was equivalent across different treatment groups (1.08 ± 0.5 e7 photons/s for 100mg/kg DSPE-liposomes versus 9.95 ± 5.2 e6 photons/s for PBS). To determine how our liposomal priming affected ctDNA performance at different tumor burdens, a similar workflow to that described for the CT26 model above was performed 1 week, 2 week, and 3 weeks after tumor inoculation. Namely, at each timepoint blood was collected retro-orbitally pre-treatment as an internal control and 1h after DSPE-liposome- or PBS-treatment. Additionally, at end point, terminal bleed blood was collected. cfDNA concentration measurement and ctDNA detection was performed on all samples as described below.

To calculate the sensitivity of the ctDNA test for tumor detection, mice were grouped as a function of tumor burden into those with small (total burden < 1.5e7 photons/s), medium (1.5e7 photons/s < total burden < 1.5e8 photons/s), and large (total burden > 1.5e8 photons/s) tumors. Subsequently, retro-orbital plasma samples were classified as ctDNA positive if the number of unique mutations detected surpassed a given mutation threshold (between 2 and 10 mutations, from lower to higher stringency of the test) and sensitivity calculated as the % of samples that were ctDNA positive per group.

### Probe panel design for ctDNA detection assay

A CT26 specific probe panel was designed by selecting 98 heterozygous SNVs from a published list (*36*). Luc-MC26 specific probe panel was designed by selecting 2000 SNVs from analyzing whole-genome sequencing (WGS) data of Luc-MC26 gDNA and BALB/c gDNA with raw depth of 30x and 15x, respectively, among which 1882 SNVs were validated by targeted sequencing.

### cfDNA extraction and quantification

Frozen plasma was thawed and centrifuged at 15,000 x g for 10 minutes to remove residual cells and debris. 1x PBS was then added into plasma to make the total volume 2.1ml for cfDNA extraction using the QIAsymphony Circulating DNA kit (cat:937556, Qiagen). The extracted cfDNA was quantified using a qPCR assay and then frozen at −20oC until ready for further processing. Plasma cfDNA concentration was quantified using a Taqman qPCR assay targeting a locus on mouse genome.

Forward primer: GGGACTCCTGCAGATCGTTA;

Reverse primer: ATCTGGCCCTATCTTCCATCCT;

Taqman probe: /56-FAM/CCTGTGGTG/ZEN/CTGAACCTATCAACAGCA/3IABkFQ/.

### gDNA extraction and shearing

Genomic DNA (gDNA) was extracted from CT26 cells, Luc-MC26 cells, or the buffy coat of the BALB/c strain using the QIAsymphony DNA Mini Kit (cat:931236, Qiagen). The extracted gDNA was sheared to 150 bp in size using a Covaris LE 220 instrument. Sheared DNA was quantified using Qubit™ dsDNA HS (cat:Q32854, ThermoFisher).

### Library construction, hybrid capture, and sequencing

Cell-free fDNA and gDNA libraries were constructed using the Kapa Hyper Prep kit (cat: 07962363001, Roche) with custom dual index duplex UMI adapters (IDT), as previously described(9). A maximum of 50μL or 20ng of extracted cfDNA or 20ng gDNA mass was used as input into library construction (LC). The prepared libraries were then quantified using the Quant-iT PicoGreen assay (cat: P11496, Invitrogen) on a Hamilton STAR-line liquid handling system. Hybrid capture (HC) using cancer cell line specific panels was performed using the xGen hybridization and wash kit (cat: 1080584, IDT) with xGen Universal blockers (cat: 1075476; IDT). For the ctDNA diagnostic test, libraries were pooled up to maximum 12-plex, with a library mass equivalent to 25 times DNA mass into LC for each sample, and 0.56 pmol/uL of a panel consisting of 120 bp long probes (IDT) targeting cancer cell line-specific mutations were applied. After the first round of HC, libraries were amplified by 16 cycles of PCR and then carried through a second HC but with half volumes of human Cot-1 DNA, xGen Universal blockers, and probes (note: we tested mouse Cot-1 DNA and did not observe any impact on the assay performance). After the second round of HC, libraries were amplified through 8-16 cycles of PCR, quantified, and pooled for sequencing (151 bp paired-end runs) with a targeted raw depth of 40,000 x per site for 20 ng DNA input. Sequencing data was processed by our duplex consensus calling pipeline as previously described, yielding measurements of the total number of mutant duplexes detected, the unique number of loci detected, and the tumor fractions detected as previously described (*12*).

### Toxicity assessment of nanoparticle priming agent

100mg/kg or 300mg/kg DSPE-liposomes (in 200ul sterile DPBS, n = 3 mice per group) or sterile DPBS (n = 3 mice per group) were injected i.v. into awake mice once a week for 3 weeks. Weight measurements were performed every other day from the first injection to 1 week after the last injection. 1 week after the last injection, terminal bleed blood was collected, and serum was used for biochemical analysis using the IDEXX Chem21 panel (performed through the Division of Comparative Medicine Laboratory at the Massachusetts Institute of Technology). Additionally, heart, lungs, liver, spleen, and kidney were harvested, fixed, paraffin-embedded, sectioned, and stained with hematoxylin and eosin and assessed by a veterinary pathologist.

### Statistical analysis

All the statistical analyses were conducted in GraphPad 9.0 (Prism). All the sample sizes and statistical tests are specified in the figure legends. For each animal experiment, groups were established before tumorigenesis or treatment with liposomes, and therefore no randomization was used in the allocation of groups. Investigators were not blinded to the groups and treatments during the experiments.

## Acknowledgments

We thank H. Fleming (MIT) for critical reading and editing of the manuscript and the Koch Institute’s Robert A. Swanson (1969) Biotechnology Center for the technical support. Specifically, we would like to thank K. Cormier and R. Bronson from the Koch Institute Histology Core for tissue processing and pathological assessment, respectively, and Dong Soo Yun from the Nanotechnology Materials Core (RRID:SCR_018674) for assistance on the TEM and cryo-TEM imaging. We would also like to thank Greg Gydush for his help setting up the data analysis pipeline leveraged for this work and Leslie Gaffney for her assistance in designing the graphics for this work.

## Author contributions

Conceptualization: J.C.L., V.A.A., S.N.B, C.M.A., S.T., K.X.

Methodology: C.M.A., S.T., K.X. T.B., S.P., Z.A., S.S., A.B., D.S., A.P.A., J.K., J.R., S.W.

Investigation: C.M.A., S.T., K.X. T.B., S.P., Z.A., S.S., A.B., D.S., A.P.A., J.K., J.R., S.W.

Visualization: C.M.A., S.T., K.X., T.B.

Funding acquisition: J.C.L., V.A.A., S.N.B, T.R.G

Project administration: C.M.A., S.T., K.X.

Supervision: J.C.L., V.A.A., S.N.B

Writing – original draft: C.M.A., S.T., K.X.

Writing – review & editing: C.M.A., S.T., K.X., A.P.A., J.C.L., V.A.A., S.N.B

## Data and materials availability

All sequencing data generated in this study will be deposited into a controlled access database.

## Supplementary Materials

Figs. S1 to S16; Data S1 to S2

