## Supplementary Material for "A nanoparticle priming agent reduces cellular uptake of cell-free DNA and enhances the sensitivity of liquid biopsies"

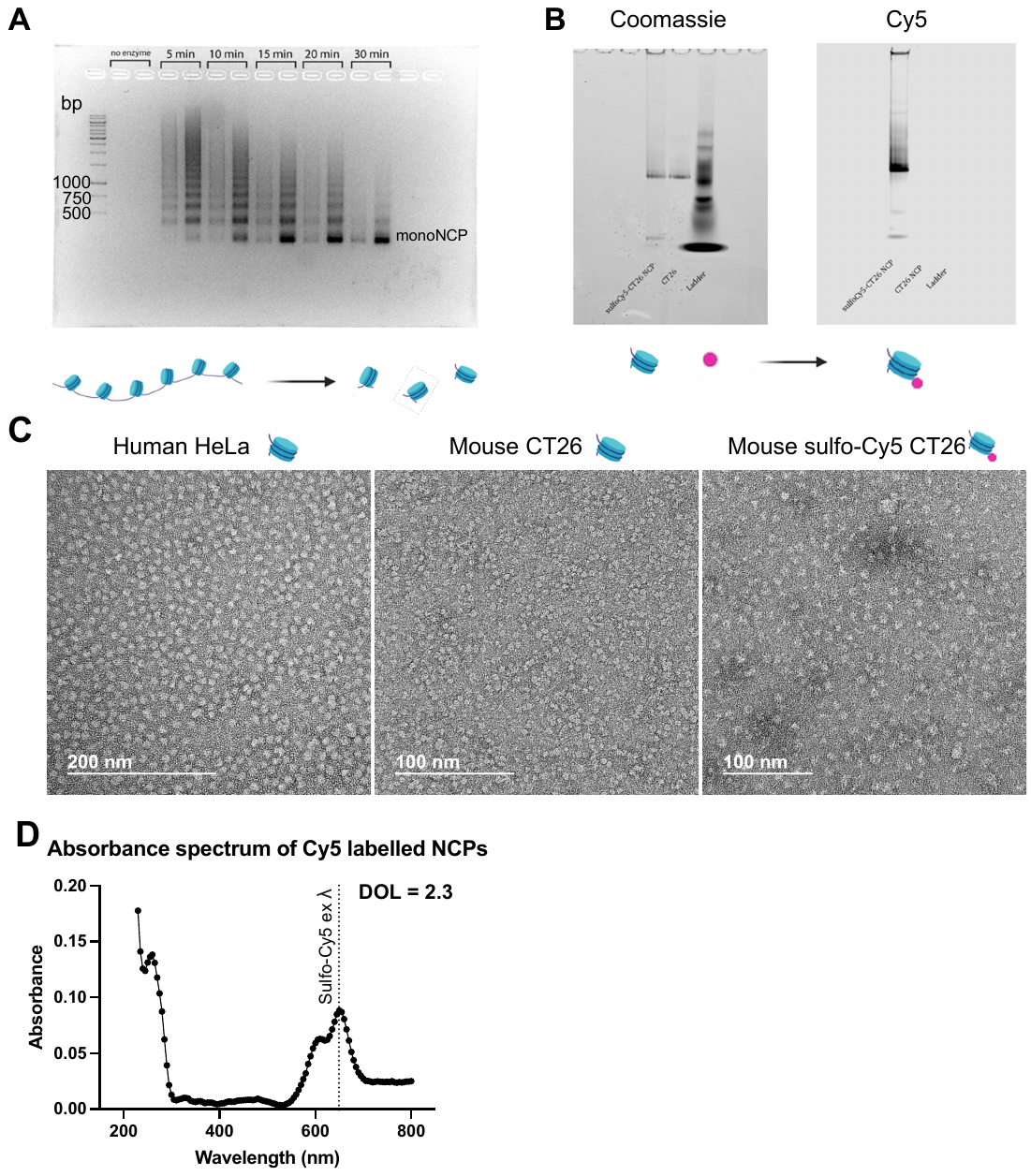

Fig. S1. Optimization of mononucleosome preparation and Cy5-labeling from the chromatin of CT26 cells. (A) Mononucleosomes (MN) were prepared by digesting the chromatin of CT26 cells for 5-30min using the “Nucleosome preparation Kit”. The MN were subjected to DNA cleanup and digestion efficiency assessed via electrophoresis through a 1.5% agarose gel. A digestion time of 30min was rendered optimal to obtain a MN preparation. (B) MN were labelled with sulfo-Cy5 dye (25 dye: 1 protein molar ratio, 4C at 550rpm overnight) and labelling confirmed by co-localization of a Cy5-positive band and a protein band in a 4-12% Tris Glycine Novex gel (100V, 2h, 4C, native running buffer). (C) Representative TEM images of unlabeled commercial human HeLa MN (left) and murine CT26 MN prepared as described in (A) prior to (middle) and after sulfo-Cy5 labeling (right). Scale bars as indicated in images. Prepared MNs remain stable after labeling and resemble those purchased commercially, albeit being in smaller in size (11.4 ± 1.3 nm (81 particles) for human versus 5.2 ± 0.7 nm (82 particles) for mouse, respectively, measured by the ImageJ software). (D) Absorbance spectrum of Cy5-labelled MNs measured using an Infinite F200 Pro reader (Tecan) shows a peak at the excitation wavelength of the sulfo-Cy5 fluorophore (649nm). Degree of labelling was estimated by calculating the protein concentration using a Nanodrop and the dye concentration using the Infinite F200 Pro reader to be 2.3 dye molecules per MN molecule.

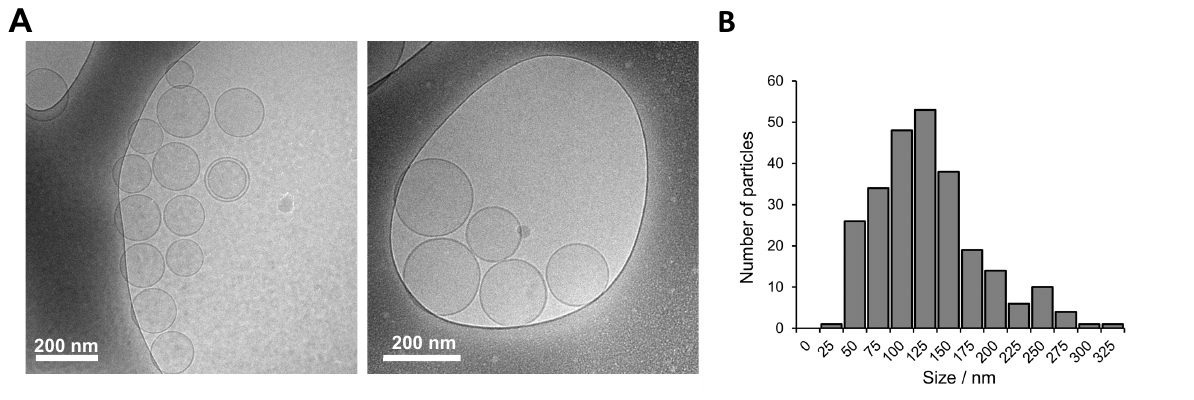

Fig. S2. Cryogenic transmission electron microscopy (cryo-TEM) images verified the vesicular morphology for DSPE-based liposomes. (A) Representative images of DSPE-based liposomes show the vesicular morphology of liposomes, with a dark opaque hydrophobic layer separating the inner hydrophilic lumen. Liposomes are mostly unilamellar. Scale bars as indicated in images. (B) Histogram of particle size was obtained by manually counting 256 particles. Each particle diameter was measured three times using the ImageJ software and the average value was recorded in the histogram.

­
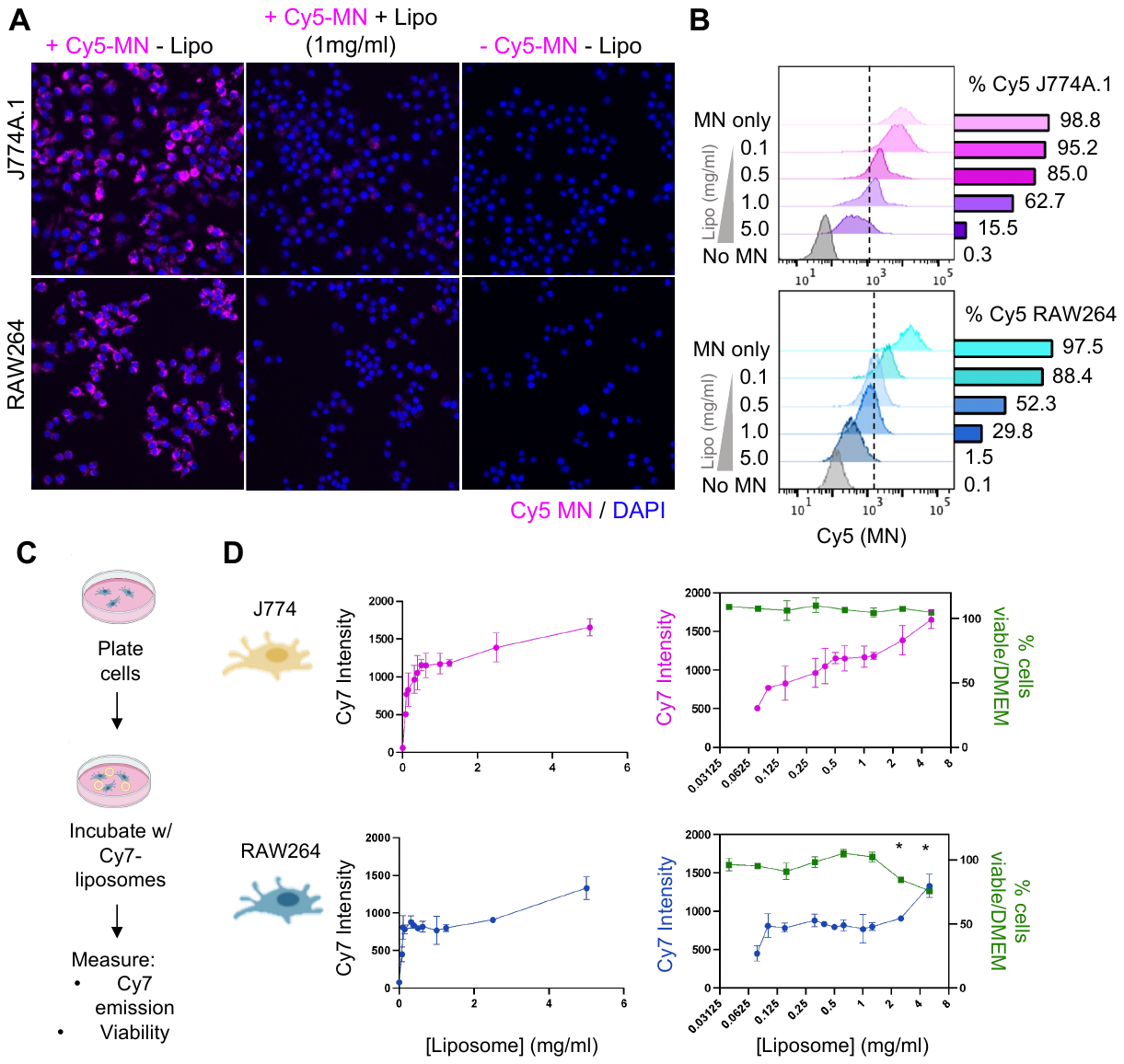

Fig. S3. DSPE liposomes inhibit the uptake of mononucleosomes in two different macrophage cell lines with minimal effects on cell viability. (A) Representative epifluorescence images and (B) flow cytometry-based quantification of Cy5-MN uptake by J774A.1 and RAW264 cells after liposome pre-treatment for 4 hours. (C) Experimental workflow to determine the viability of cells 4h post-liposome treatment and to characterize the uptake of Cy7-liposomes. (D) Overlay of cell viability measurements and Cy7-liposome measurements at a range of liposome concentrations. n = 3 wells per condition for A-D, * P < 0.05, two-tailed ANOVA.

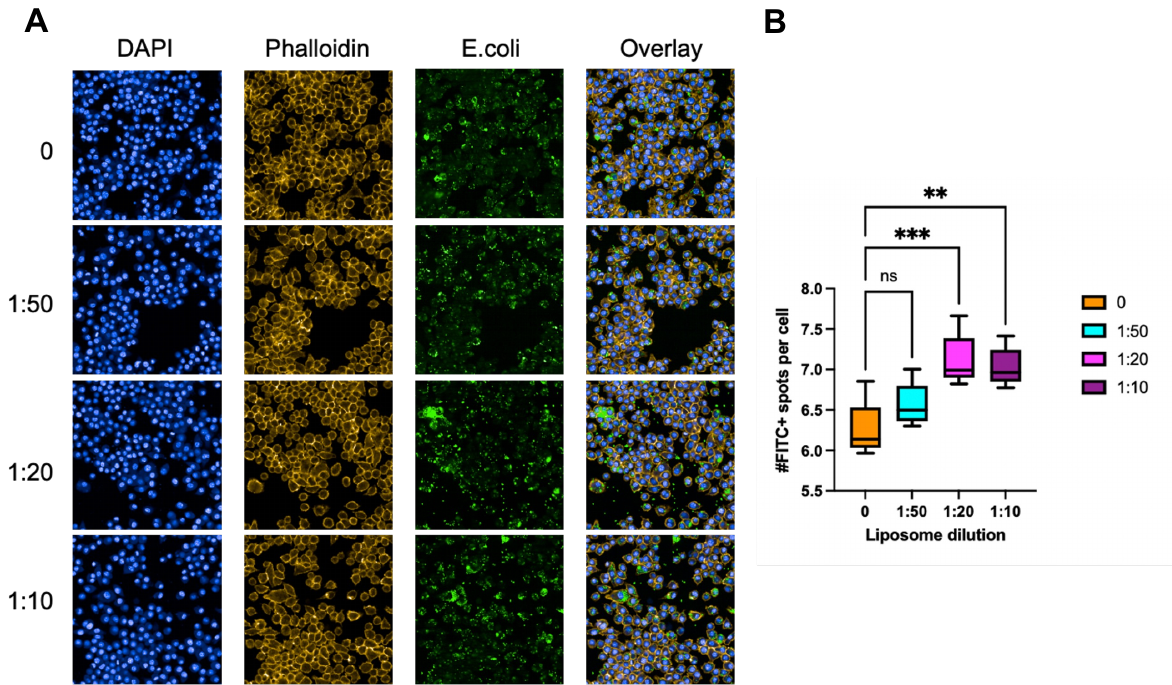

Fig. S4. DSPE-liposomes do not inhibit phagocytosis of E.Coli particles in J774A.1 cells. (A) Representative epifluorescence images and (B) image analysis-based quantification of E.Coli uptake by J774A.1 cells after liposome pre-treatment for 1-hour.

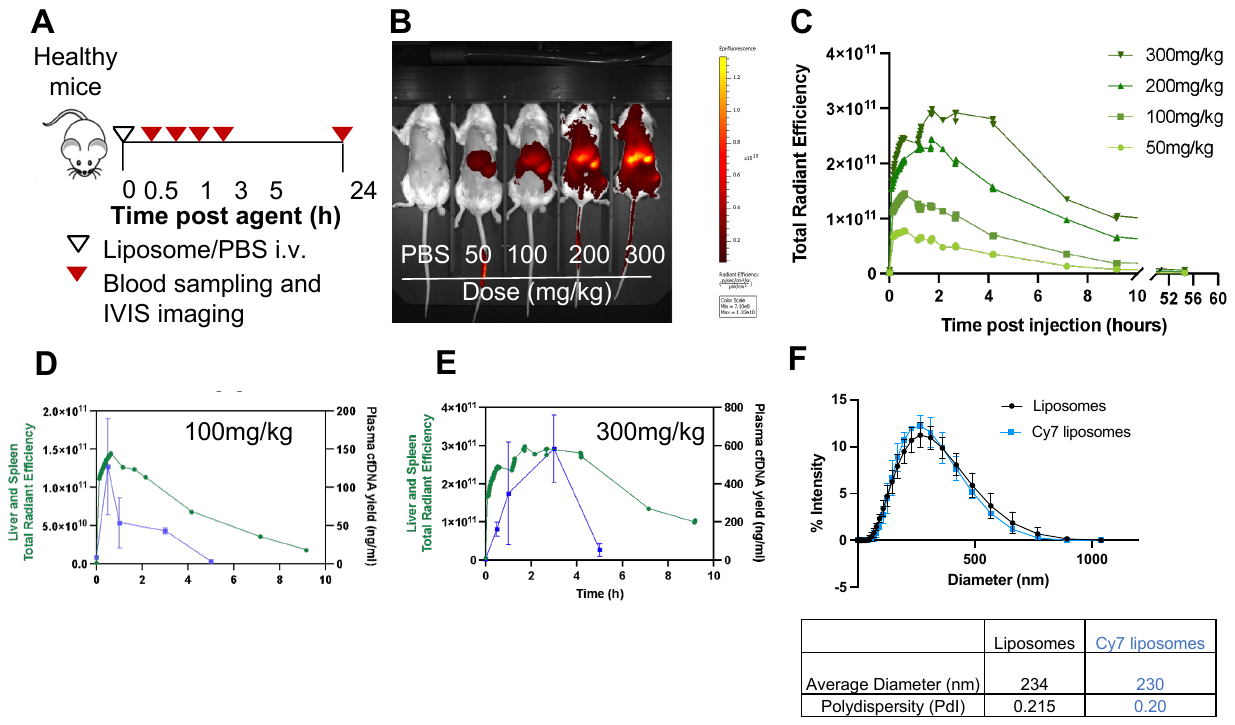
**Fig. S5.** ***In vivo* in healthy mice liposomes occupy the two major organs of the MPS.** (**A**) Experimental timeline to investigate the kinetics of liposome biodistribution and of cfDNA concentration changes in the plasma of healthy Balb/c mice following liposome administration. (**B**) Image of liposome accumulation in the liver and spleen of mice 10 min after administration. (**C**) IVIS-based quantification of liposome accumulation in the liver and spleen (as defined by a region of interest occupying the upper abdominal area of mice that overlaps with the anatomical positioning of the liver and the spleen) over time. Overlay of cfDNA plasma levels and spleen and liver accumulation after dosing liposomes at (**D**) 100 mg/kg or (**E**) 300 mg/kg showing that the maximum accumulation of liposomes in target organs is achieved prior to the peak in plasma cfDNA levels. This suggests that accumulation in MPS organs may be driving the observed trends in plasma cfDNA concentrations. (**F**) DLS-based characterization of non-fluorescent and Cy7-liposomes.

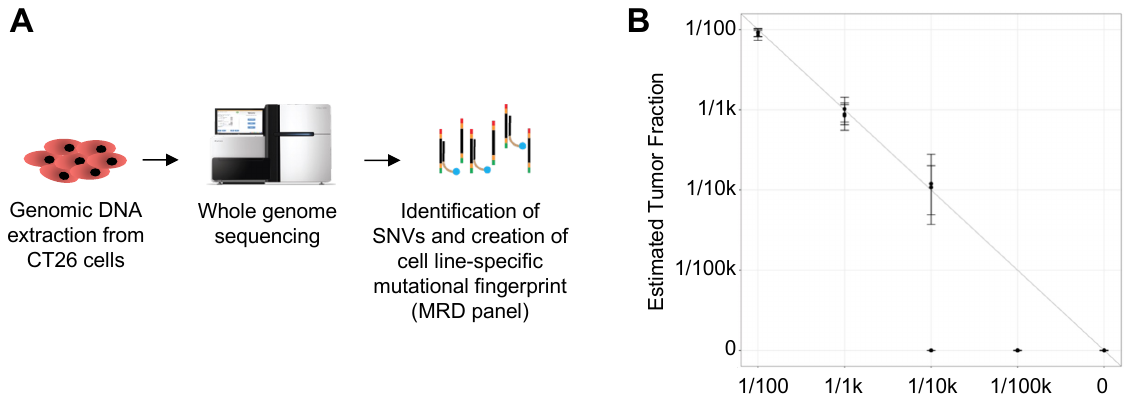

**Fig. S6. Design of a ctDNA test for the CT26 bi-flank tumor model.** (**A**) Schematic for the design of a CT26-specific tumor fingerprint panel. (**B**) Validation of a ctDNA test tracking 98 mutations of the CT26 bi-flank mouse model. The assay estimated tumor fractions are concordant with the actual tumor fractions down to 1:10k. Samples were prepared by serially diluting CT26 tumor gDNA into normal gDNA extracted from the buffy coat of healthy mice (n=3 biological replicates).

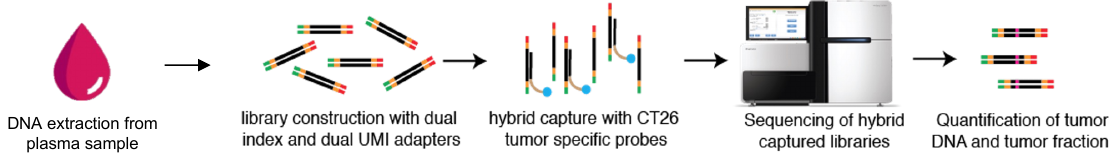

Fig. S7. Schematic for the processing of plasma samples to detect mutations. DNA is extracted from plasma samples, libraries are constructed using the Kapa Hyper Prep kit (Roche) and custom dual UMI adapters (IDT), regions of interest on the genome are captured with CT26 tumor specific probes and sequencing is performed. Following alignment and duplex consensus sequences assembly, the numbers of mutant and WT molecules are quantified, and tumor fractions are calculated using previously published methods (*12*).

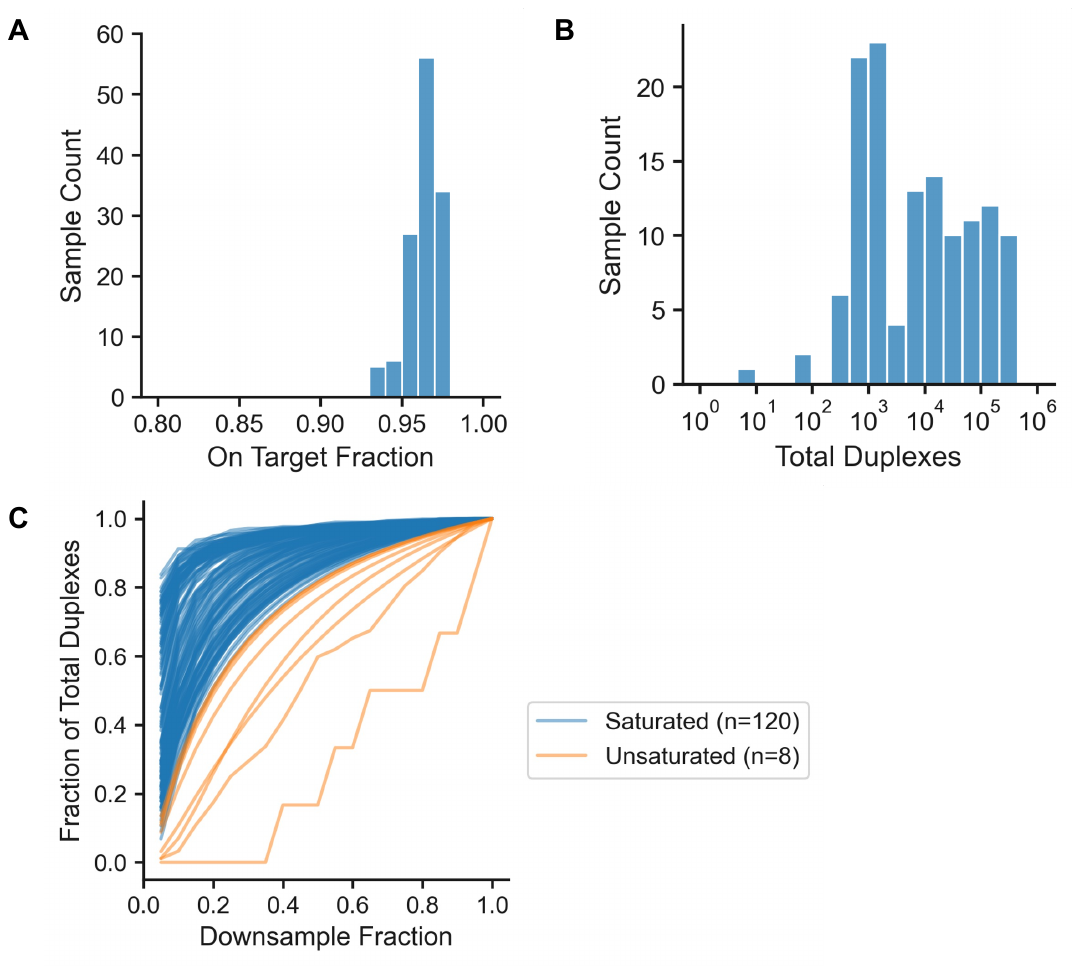

Fig. S8. Technical performance of the ctDNA diagnostic test tracking 98 mutations in the CT26 bi-flank tumor model. Histograms of (A) the on-target rate and (B) the total number of duplexes recovered. (C) Fraction of total duplexes after down sampling raw reads. We considered a sample to be saturated if it still contained at least 95% of its duplexes after down sampling to 80% of its raw reads.

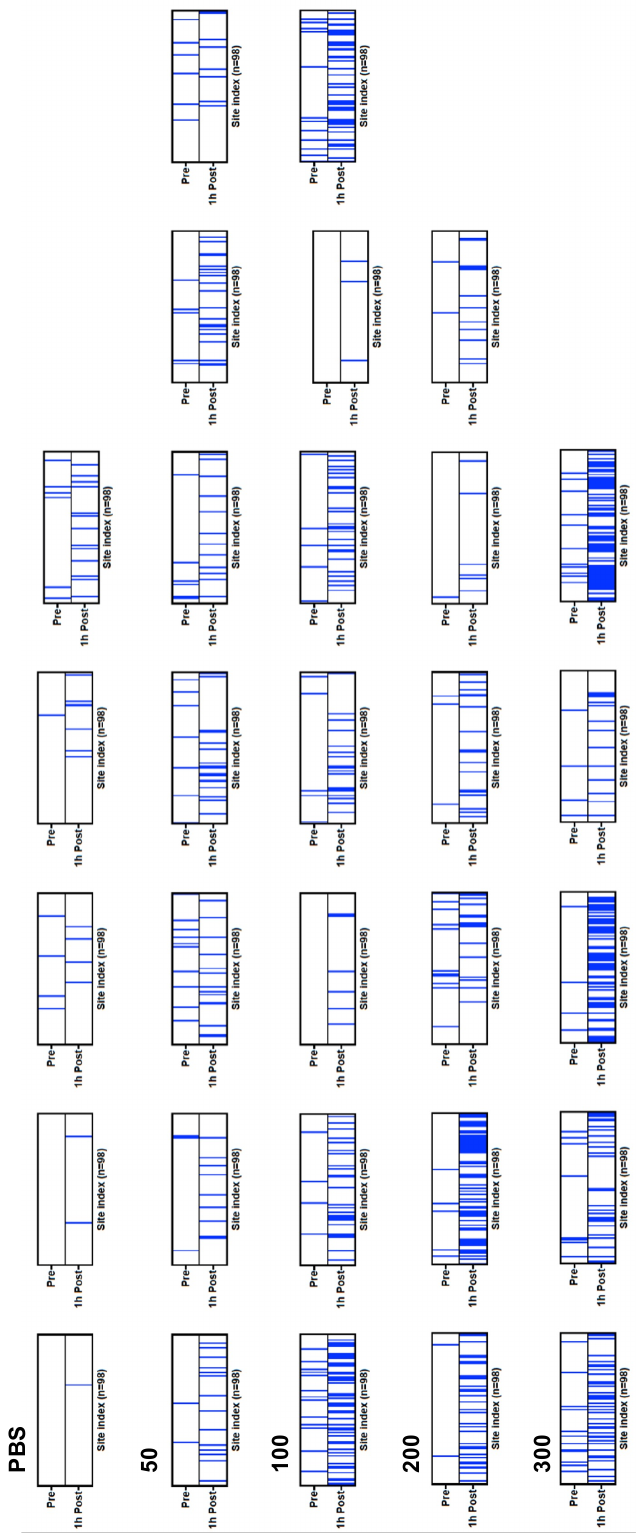

**Fig S9. Heatmap representation of the mutational fingerprints detected pre-treatment and 1 hour post-treatment for each mouse for the CT26 bi-flank tumor-bearing mouse cohort**. The top row represents mutations detected pre-treatment and the bottom row mutations detected 1h post-treatment in a given plasma sample. Each vertical band corresponds to a different site in our 98-site mutational panel and is colored blue if detected at least once in the plasma sample. 4 representative heatmaps for each group were selected to show in Fig 4e. Each row represents a different treatment group.

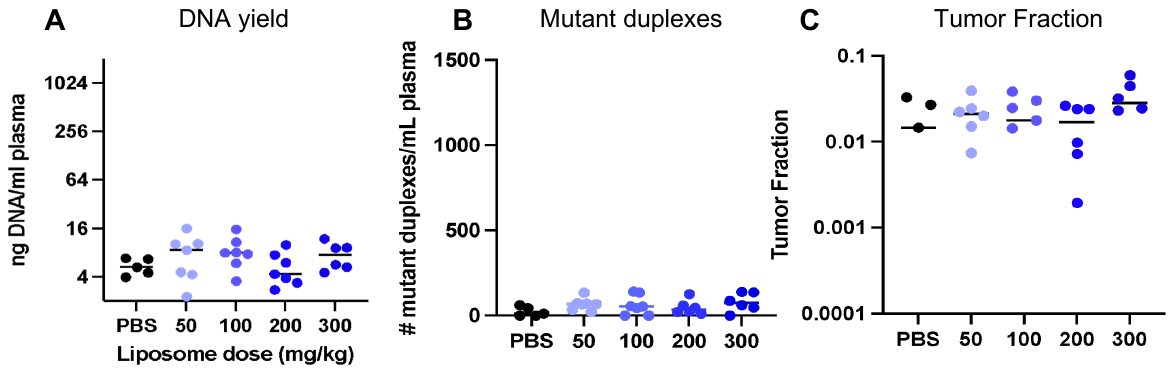
 Fig S10. (A) Pre-treatment cfDNA yields, (B) mutant duplexes recovered and (C) tumor fractions are comparable across groups in the flank tumor-bearing mice cohort. We observed no statistical differences between the pre-treatment cfDNA yields of PBS-treated or liposome-treated mice at all doses. Mann-Whitney, two-tailed test. PBS vs. 50mg/kg p=0.4318, P BS vs. 100mg/kg p=0.1061, PBS vs. 200mg/kg p=0.6389, PBS vs. 300mg/kg p=0.1775.

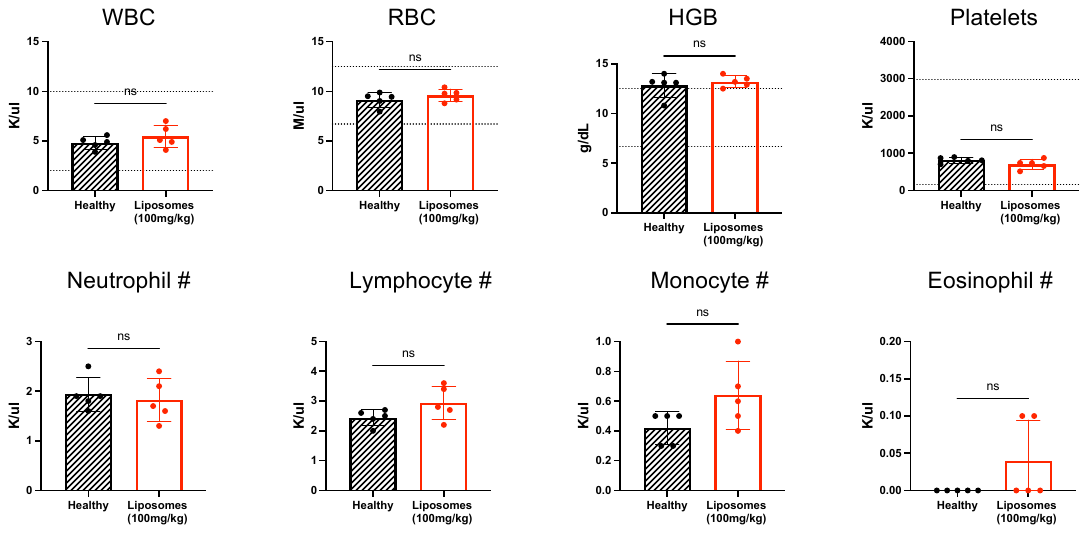

Fig S11. Liposome treatment does not affect the amount of different cellular subtypes in blood. Absolute amounts of different cellular subtypes in the blood 1 hour after liposome administration. Healthy mice were treated with 100mg/kg liposomes or PBS and blood drawn retro-orbitally 1 hour after administration for complete blood count assessment. Dotted lines represent reference ranges (n=5 per group, ns = P > 0.05, two-tailed unpaired t-test).

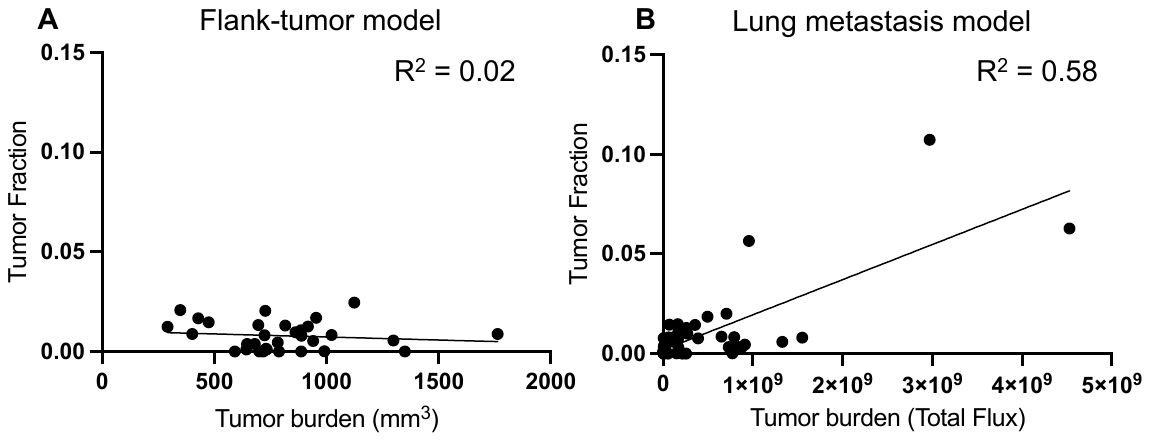

**Fig S12. Scatterplot of pre-treatment tumor fraction versus tumor burden shows higher correlation in (B) MC26 lung metastasis model than in (A) CT26 bi-flank tumor model,** suggesting earlier and more consistent access to the vasculature. Higher tumor fractions also demonstrate greater access of ctDNA into circulation.

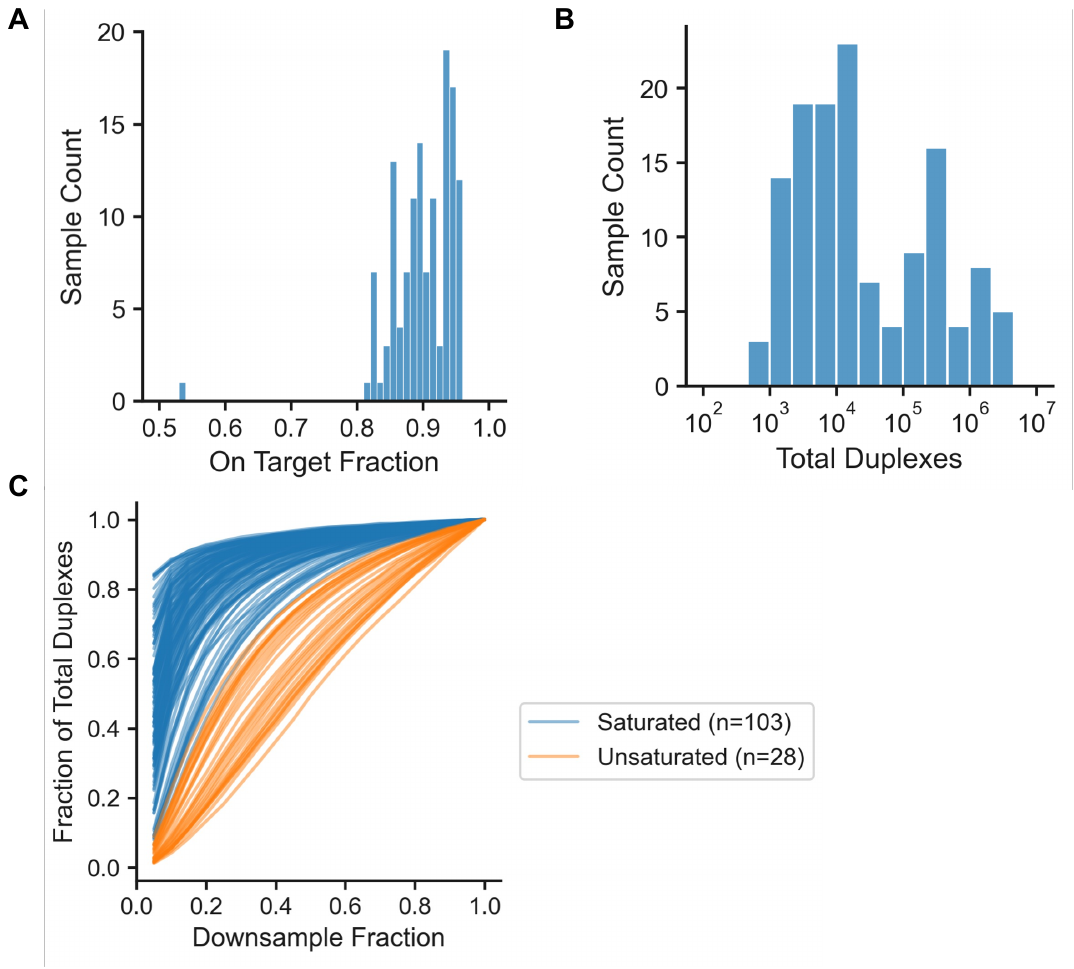

**Fig. S13. Technical performance of the ctDNA diagnostic test tracking 1822 mutations of the Luc-MC26 lung metastasis model.** Histograms of (**A**) the on-target rate and (**B**) the total number of duplexes recovered. (**C**) Fraction of total duplexes after down sampling raw reads. We considered a sample to be saturated if it still contained at least 95% of its duplexes after down sampling to 80% of its raw reads.

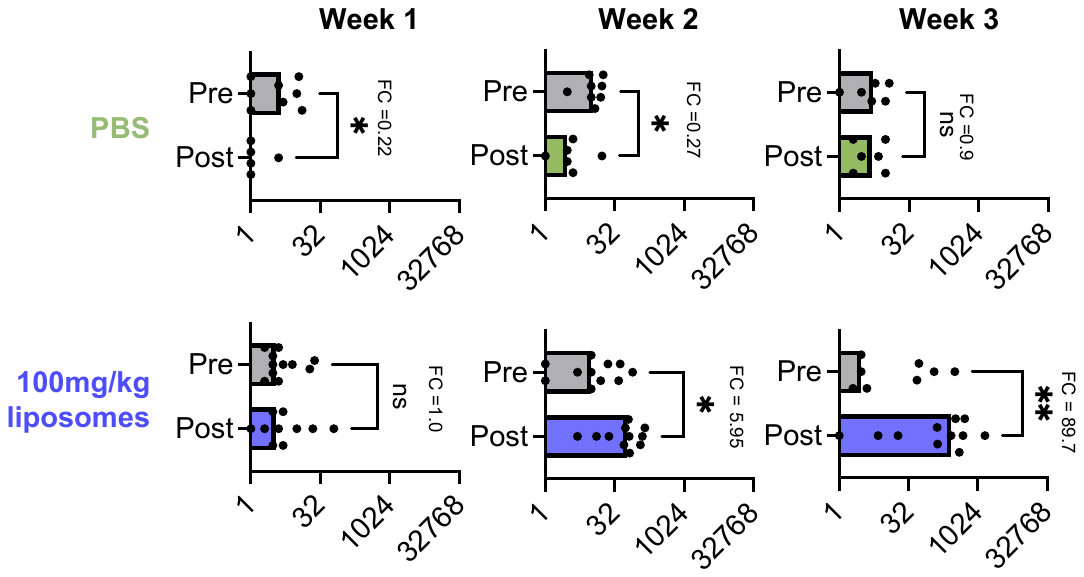

**Fig. S14. Number of unique mutational sites detected in Luc-MC26 tumor bearing mice increases with liposome treatment.** Blood was drawn prior to i.v. administration of PBS or liposomes (100mg/kg) and 1 hour after treatment 1 week, 2 weeks and 3 weeks after tumor inoculation. Liposome group had n=11-12 mice and PBS group had n=6-8 mice. Fold changes (FC) in the number of unique sites detected are shown (1822 total sites). Liposome treatment resulted in significant increases in the number of sites detected post treatment at weeks 2 (5.95-fold) and week 3 (89.7-fold), whereas PBS treatment did not increase the number of sites detected. ns P > 0.05, * P < 0.05, ** P <0.01, two-tailed Mann-Whitney test.

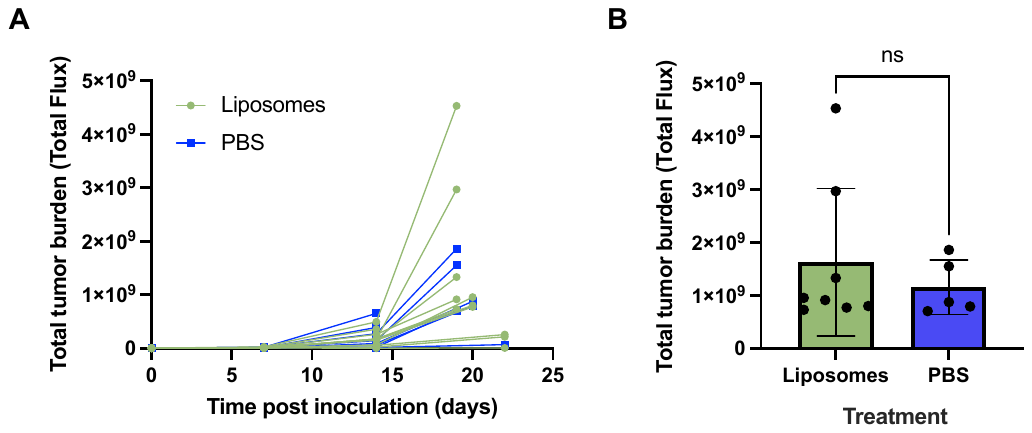

**Fig. S15. *In vivo* luminescence imaging of tumor burden over time shows no difference in tumor growth between liposome and PBS-treated mice.** Luc-MC26 tumors were inoculated via tail vein and IVIS imaging performed once a week during tumor progression. Mice were treated 3 times with liposomes (100mg/kg) or PBS; on days 6, 14 and between days 19 and 21. **(A)** Total tumor burden over time and **(B)** tumor burden at days 19 and 20 post-tumor inoculation for each treatment group. ns P > 0.05, two-tailed unpaired t-test.

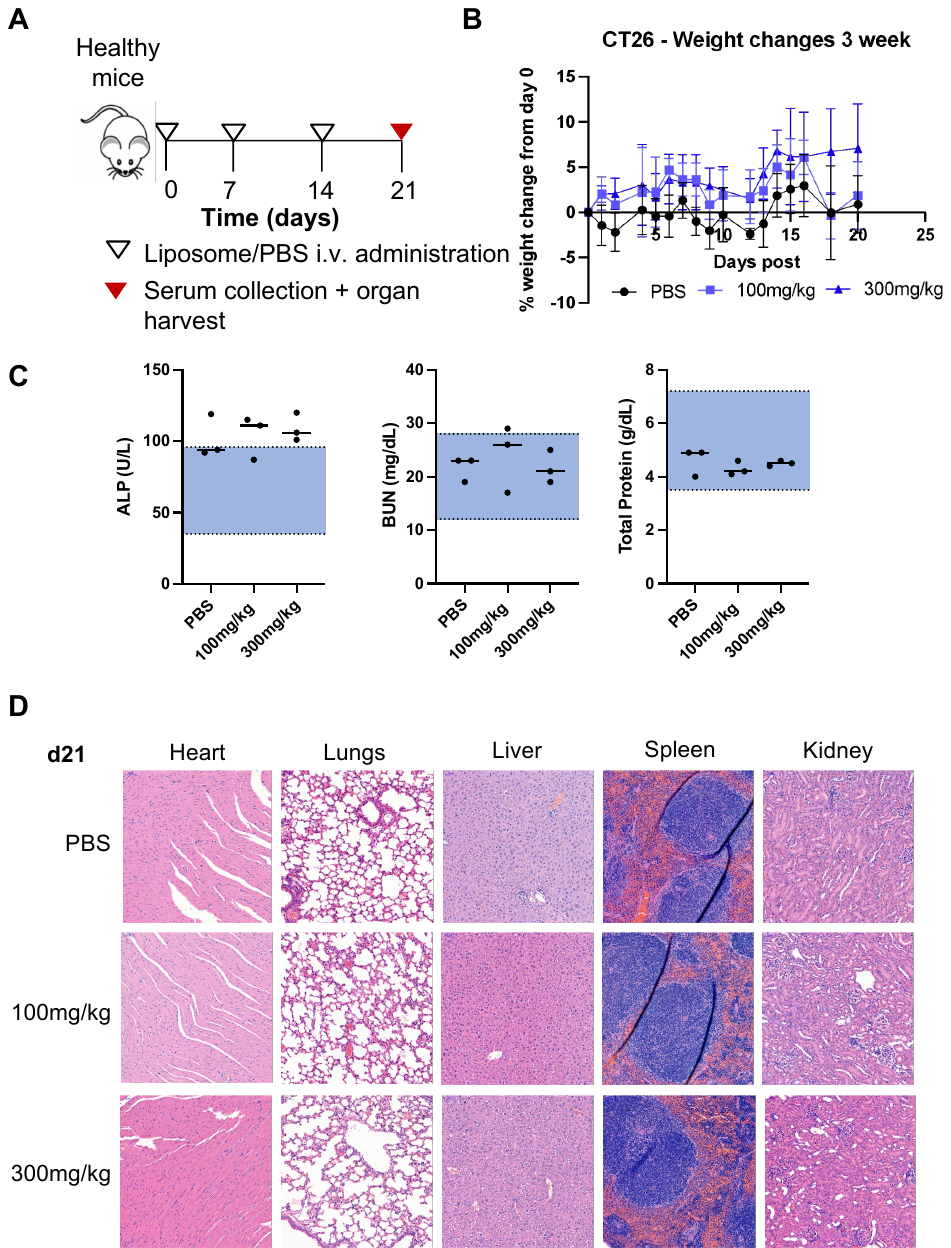

**Fig 16. Liposomal contrast agent incurs no signs of acute toxicity or weight loss after 3 doses.** (**A**) Experimental timeline to assess the toxicity of the priming agent in mice. (**B**) Liposomes at either dose incurred no significant weight loss over the treatment course. (**C**) Serum chemistry assessing a basic liver and kidney panel showed no significant difference between PBS- and liposome-treated mice. (**D**) 7-days after the third liposome dose, organs (heart, lung, liver, spleen, and kidney) were collected, fixed, embedded in paraffin, and stained with hematoxylin & eosin. Analysis by a veterinary pathologist confirmed that tissues from liposome-injected mice appeared similar to PBS injected controls, exhibiting no signs of toxicity. Study was done with n = 3 mice per group and images from representative animals are shown.

**­­­Data S1. (separate file)**

List of 98 tracked sites in CT26 panel, each carrying a mutation specific to the CT26 cell-line.

Data S2. (separate file)

List of 1,822 tracked sites in Luc-MC26 panel, each carrying a mutation specific to the Luc-MC26 cell-line.
